# Yeast cell responses and survival during periodic osmotic stress are controlled by glucose availability

**DOI:** 10.1101/2023.02.17.528926

**Authors:** Fabien Duveau, Céline Cordier, Lionel Chiron, Matthias LeBec, Sylvain Pouzet, Julie Séguin, Artémis Llamosi, B. Sorre, Jean-Marc Di Meglio, Pascal Hersen

## Abstract

Natural environments of living organisms are often dynamic and multifactorial, with multiple parameters fluctuating over time. To better understand how cells respond to dynamically interacting factors, we quantified the effects of dual fluctuations of osmotic stress and glucose deprivation on yeast cells using microfluidics and time-lapse microscopy. Strikingly, we observed that cell proliferation, survival and signaling depend on the phasing of the two periodic stresses. Cells divided faster, survived longer and showed decreased transcriptional response when fluctuations of hyperosmotic stress and glucose deprivation occurred in phase than when the two stresses occurred alternatively. Therefore, glucose availability regulates yeast responses to dynamic osmotic stress, showcasing the key role of metabolic fluctuations in cellular responses to dynamic stress. We also found that mutants with impaired osmotic stress response were better adapted to alternating stresses than wild-type cells, showing that genetic mechanisms of adaptation to a persistent stress factor can be detrimental under dynamically interacting conditions.

## Introduction

Cells have evolved to survive in a broad range of environmental conditions with multiple factors (e.g., temperature, nutrients, light, humidity, pathogens…) varying in space and time. They can monitor their environment and constantly adapt their physiology to stress caused by environmental fluctuations. Experiments in which cells are dynamically probed with time-varying stress signals are required to obtain a quantitative understanding of how signaling pathways and gene regulatory networks confer cellular adaptability to environmental changes^1^. The development of microfluidics systems to study the frequency responses of cellular functions^2–5^ (*e.g.,* signaling pathways, gene regulatory networks) has been instrumental in the adoption of the concepts of dynamic systems and information processing in biology. More recent methodological developments in the field of control theory have enabled time-varying perturbations to be used to control cellular gene expression or signaling pathways via computer-based external feedback loops^6–11^. In short, methods are now mature to study cells as dynamical systems.

Most studies of cellular stress responses have focused on a single environmental stress in an otherwise maintained environment. However, how cells respond to stress often depends on the interaction between several environmental factors. For instance, changing the metabolic environment (*e.g.,* carbon source type and concentration) can profoundly affect cell physiology (*e.g.,* respiration and fermentation in yeast) and alter stress responses^12,13^. More generally, resource allocation (*i.e*., how cellular resources are shared between several cellular functions) is an important fundamental^14,15^ (*e.g.,* understanding growth laws) and applied topic^16^ (*e.g.,* design of robust synthetic gene circuits and bioproduction). Specifically, cells face decision-making problems when exposed to stress and to variation in their metabolic environment. Routing resources towards stress response mechanisms may deprive other important processes (*e.g.,* cellular maintenance, proliferation) and decrease competitive fitness in an environment periodically scarce of metabolic resources. Conversely, routing resources towards cell proliferation may reduce survival in stressful conditions and therefore also reduce fitness. The tradeoff between proliferation and stress responses can be an important determinant of cell fitness in a dynamic environment^14,15,17,18^. Yet, the extent to which what is known in rich, constant metabolic conditions, remains valid under low or fluctuating nutrient availability remains an open question. Here, we address this broad question by studying the synergistic and antagonistic effects of time-varying osmotic stress and glucose deprivation on the growth of budding yeast cells.

The adaptation to hyperosmotic stress in the budding yeast *S. cerevisiae* involves an adaptive pathway—the HOG pathway—that has been extensively described at the molecular level^19^ as well as biophysical and integrative levels through mathematical and computational models^20–26^. Quantitative descriptions of the dynamics of osmotic stress responses were achieved using microfluidics to generate time-varying perturbation of the osmolarity of the environment while observing signaling activity and the transcriptional responses of key players in the HOG pathway at the single-cell resolution via time-lapse microscopy^4,5,27^.

When external osmolarity increases, accumulation of intracellular glycerol is required to restore the cellular osmotic balance^28^. At the molecular level, osmotic stress signaling is orchestrated by a mitogen activated protein kinase (MAPK) cascade, which culminates in double phosphorylation and nuclear accumulation of the MAPK protein Hog1p and differential regulation of hundreds of genes^29,30^. In particular, GPD1 (NAD-dependent glycerol-3-phosphate dehydrogenase), a key enzyme involved in production of glycerol from glucose, is upregulated after hyperosmotic stress (**Figure 1a**). Phosphorylated Hog1p also triggers several processes in the cytoplasm that are essential for osmoregulation^21,22^, including cell-cycle arrest^31–33^. Dynamically, the HOG pathway behaves as a low pass filter that drives (perfect) adaptation through at least two layers of feedback loops that allow for deactivation of the pathway^21^ (transcriptionally and within the cytoplasm). Notably, the HOG pathway can be hyper-activated when stimulated at high frequencies, which drastically slows down the cell cycle^27^. Although very informative—and an excellent example of how biological and physical concepts can be combined to obtain a comprehensive description of gene regulatory network dynamics—these studies were carried out in a glucose-rich environment, which insulates metabolic needs from osmotic stress adaptation requirements. Glucose is not only needed for growth, but also for production of glycerol and the transcriptional feedback loop that deactivates the HOG pathway^12,21^; thus, cells may employ a decision mechanism to share glucose internally between these processes, particularly when glucose is scarce, or its availability fluctuates. **More generally, despite the known importance of the metabolic state in cellular adaptation to stress, the systemic interactions between cellular maintenance, growth and stress responses remain unexplored.**

**Caption Figure 1.**
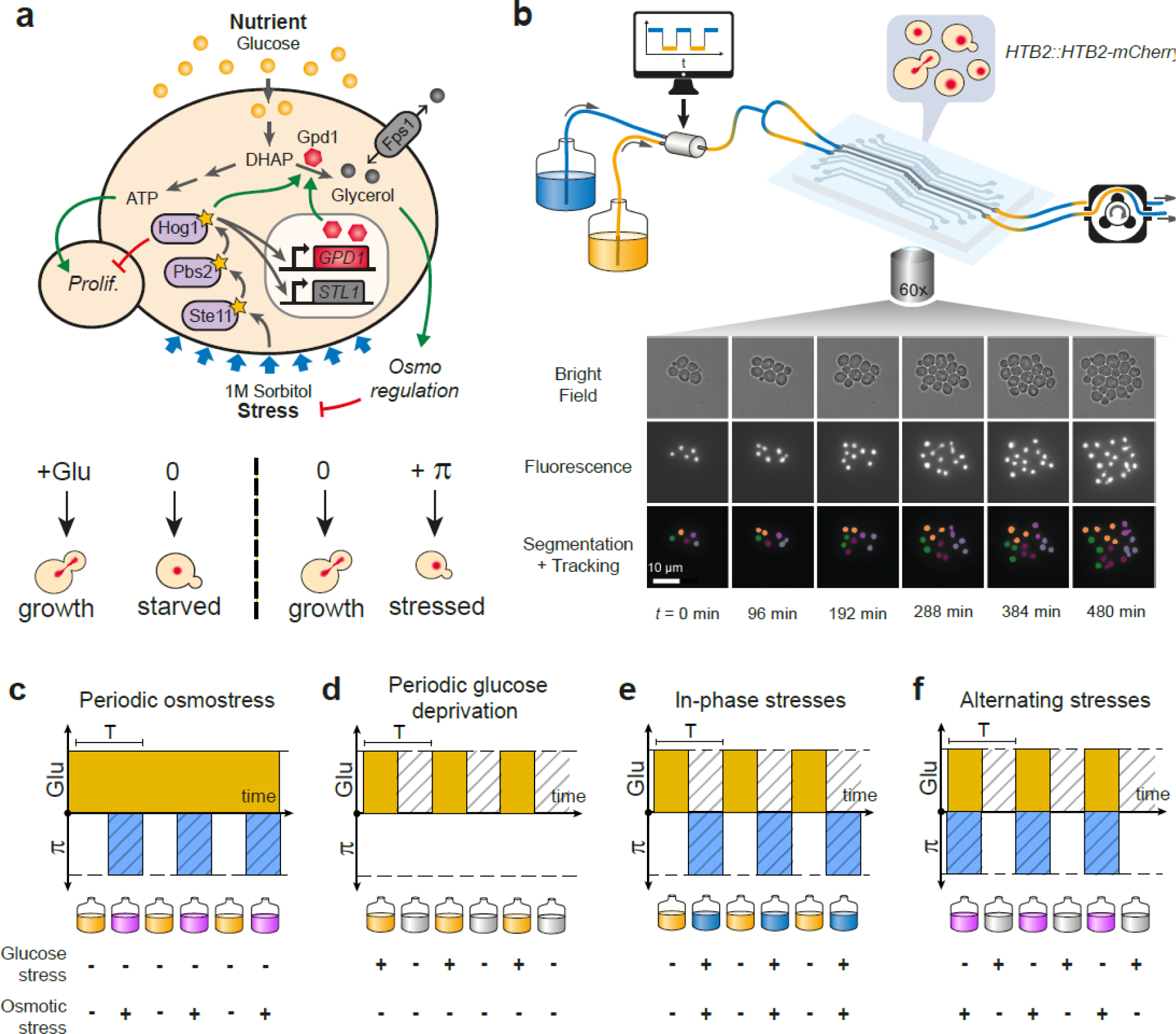
Live imaging of yeast cells grown in periodically fluctuating environments. **(a)** Overview of the hyperosmotic stress response in yeast. Both glucose deprivation and osmotic stress lead to cell cycle arrest—through different molecular mechanisms. Yeast cells maintain osmotic equilibrium by regulating the intracellular concentration of glycerol. Glycerol synthesis is regulated by the activity of the HOG MAP kinase cascade that acts both in the cytoplasm (fast response) and on the transcription of target genes in the nucleus (long-term response). For simplicity, we only represented on the figure genes and proteins involved in this study. **(b)** Sketch of the microfluidic setup used to generate a time-varying environment and achieve timelapse imaging of yeast cells. Bright field and fluorescence images are captured every 6 minutes at 25 positions for 12‒24 hours depending on the experiment. Nuclei expressing HTB2-mCherry fusion protein are segmented and tracked over time to compute the cell division rate as a function of time. (**c-f)** The four periodically varying environments used in this study. **(c)** Periodic osmotic stress: Cells are periodically exposed to hyperosmotic stress (1 M sorbitol) in a constant glucose environment (2% or 0.1%). **(d)** Periodic glucose deprivation: environment alternates between presence and absence of glucose. **(e)** In-phase stresses (IPS): periodic exposure to glucose in absence of hyperosmotic stress followed by glucose depletion with hyperosmotic stress (1 M sorbitol). **(f)** Alternating stress (AS): periodic exposure to glucose with hyperosmotic stress (1 M sorbitol), followed by glucose depletion without hyperosmotic stress. (**c-f)**. Hatching represents stress, blue indicates presence of sorbitol; orange, presence of glucose.

We address this question by monitoring the growth of yeast cells subjected to periodic variations in both osmolarity and glucose availability. To determine how resource allocation impacted cell growth, we compared two regimes of dual fluctuations that differed in the phasing of hyperosmotic stress and glucose deprivation. We showed that cell division rates, death rates and biological responses at the signaling and transcription levels are different when cells are exposed simultaneously (in-phase stresses) or alternatively (alternating stresses) to glucose deprivation and hyperosmotic stress. Therefore, yeast responses to osmotic stress are regulated by the presence of external glucose, indicating that the metabolic environment is a key factor when quantitatively assessing stress response dynamics. More globally, our study suggests that applying dual periodic perturbations is a powerful method to probe cellular dynamics at the system level and, more specifically, to clarify the role of the metabolic environment in the dynamics of cellular decision making.

## Results

### A microfluidic system to study the interaction between two environmental dynamics

We used a custom microfluidic device to monitor the growth of yeast cells exposed to periodic environmental fluctuations for up to 24 hours. Cells were imaged every 6 minutes in microfluidic chips containing five independent sets of channels connected to five growth chambers (**Figure 1b**; **Figure 1 – figure supplement 1**), allowing five different conditions per experimental run, with five technical replicates for each condition. Computer-controlled fluidic valves were programmed to generate temporal fluctuations of the media dispensed to cells with rapid transitions (< 2 minutes) from one medium to another (**Figure 1 – figure supplement 1**). The rate of cell division was then quantified using automated image analyses (see Methods). With this experimental system it is possible to determine not only how temporal fluctuations of individual parameters of the environment (e.g., a repeated stress or carbon source fluctuations) impact cell proliferation but also what are the impacts of the dynamic interactions of two environmental parameters. Here, we specifically study how periodic fluctuations of a metabolic resource (glucose concentration switching between 0% and either 2% or 0.1% w/v) and osmotic stress (sorbitol concentration switching between 0 M and 1 M) interact to alter the proliferation of yeast cells (**Figure 1**).

### Division rate correlates negatively with the frequency of osmotic stress but positively with the frequency of glucose availability

To determine how the temporal dynamics of osmotic stress altered cell proliferation, we first measured the division rate of yeast cells exposed to fluctuations between 1 M sorbitol and no sorbitol at periods ranging from 12 minutes to 480 minutes. In these experiments the time-averaged osmotic concentration was constant (i.e., cells were exposed to 1 M sorbitol half of the time in all conditions), which is important when studying the effects of the *frequenc*y, and not *intensity,* of osmotic stress on cell dynamics. The average division rate strongly decreased as the frequency of osmotic shock increased (**Figure 2a**), both in 2% glucose (2.2-fold reduction of division rate between periods of fluctuation T = 192 minutes and T = 12 minutes) and in 0.1% glucose (3.8-fold reduction of division rate between periods of fluctuation T = 192 minutes and T = 12 minutes). These results are consistent with findings from W. Lim and colleagues^27^ who attributed the drastic decrease in cellular growth observed at high frequency of osmotic shocks to overactivation of the HOG pathway. However, we also observed a clear negative relationship between the frequency of hyperosmotic stress and the division rate of HOG pathway mutants (**Figure 2 – figure supplement 3d-f**), indicating that the growth slowdown was not only explained by overactivation of the HOG pathway. The temporary reduction of division rate observed in response to a hyperosmotic shock in wild-type (**Figure 2e-f**; **Figure 2 – figure supplement 1; Figure 2 – figure supplement 2a-c)** and mutant **(Figure 2 – figure supplement 3b,c**) cells could also contribute to the negative relationship between division rate and frequency of osmotic stress. This negative relationship and the fact that cellular growth can rapidly recover after exposure to high osmolarity (**Figure 2e-f**) both indicate that yeast cells are more sensitive to *repeated* than *persistent* hyperosmotic stress.

**Caption Figure 2.**
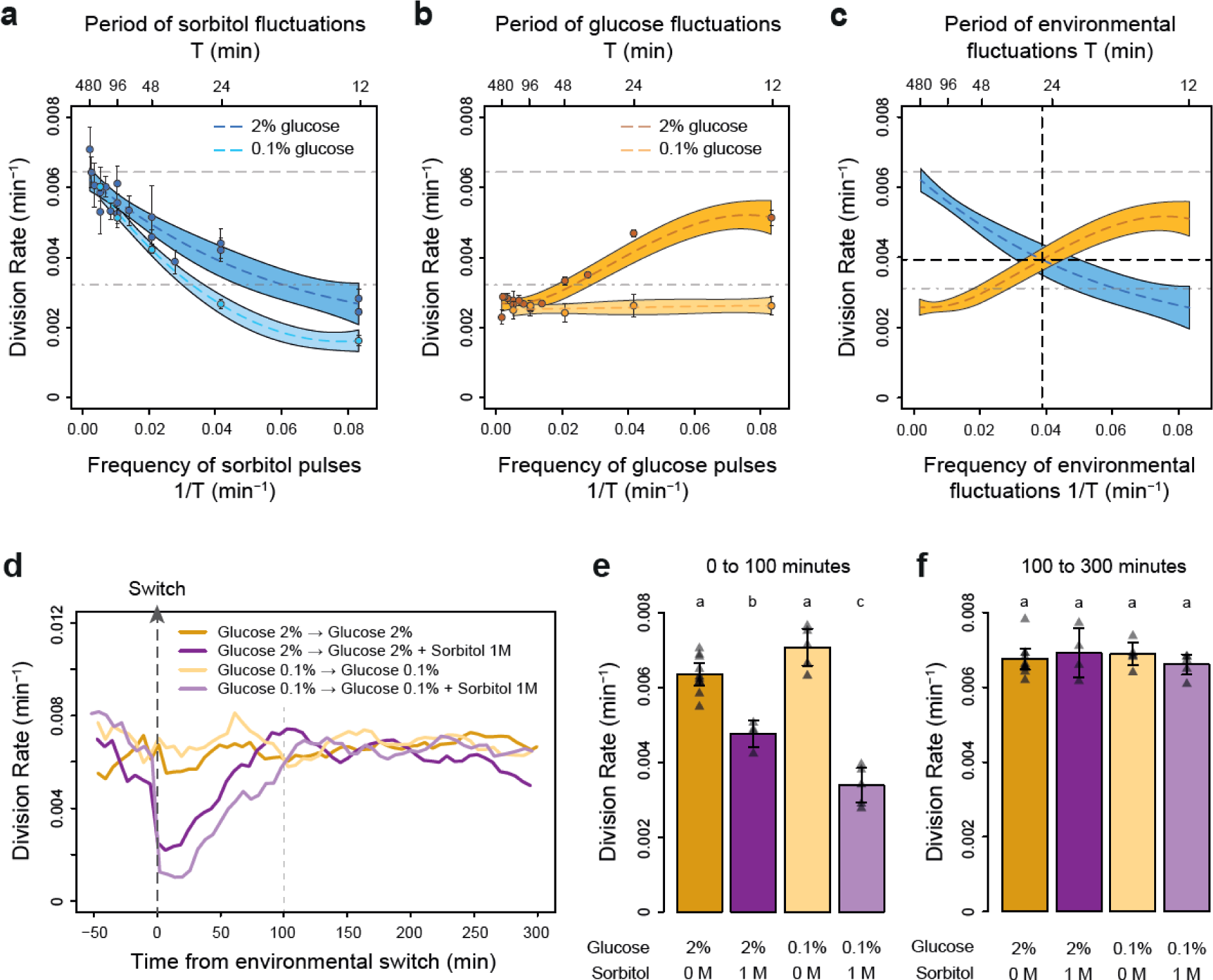
The frequencies of osmostress and glucose availability impact cell division rates in opposite ways. **(a,b)** Impact of the frequency of periodic osmotic stress **(a)** and glucose deprivation **(b)** on the average division rate. Each dot shows the mean division rate measured in different growth chambers of the microfluidic chip. Error bars are 95% confidence intervals of the mean. Colored dotted lines are Loess regressions obtained using a smoothing parameter of 0.66. Colored areas represent 95% confidence intervals of the regression estimates. Gray dashed lines show the average division rate in the absence of sorbitol (no osmotic stress) in 2% glucose (top line) and dash-dotted lines show half this average division rate (bottom line). **(c)** Overlay of the Loess regressions shown in **(a)** and **(b)** at 2% glucose. The frequency and division rate at which the two regression curves intersect are highlighted by vertical and horizontal black dotted lines. **(d)** Temporal dynamics of the division rate of cells exposed to sustained hyperosmotic stress (purple lines, 1 M sorbitol added at *t* = 0 min) or to standard conditions (orange lines) with 2% glucose (darker lines) or 0.1% glucose (lighter lines). Each curve represents the “instantaneous” division rate calculated every 6 minutes across sliding windows of 36 minutes (see Methods) and averaged for cells imaged at several positions in the microfluidic chip. **(e,f)** Cell division rates measured **(e)** from 0‒100 minutes and **(f)** from 100‒300 minutes after addition of 1 M sorbitol. Triangles represent the average division rate measured in different growth chambers of the microfluidic chips. Error bars are 95% confidence intervals of the mean division rate among growth chambers. The initial number of cells analyzed among replicates ranged from 75 to 372 in **(a)**, from 58 to 342 in **(b)** and from 220 to 776 in **(d-f)**.

Next, we wondered whether the frequency of a different type of environmental fluctuation would also affect cell division rate. To answer this question, we quantified the division rate of cells exposed to periodic transitions between a medium without carbon source and the same medium complemented with either 0.1% or 2% glucose at periods ranging from 12 to 480 minutes. In contrast to the negative effect of osmotic stress frequency, we observed a positive relationship between the frequency of glucose availability and division rate (**Figure 2b**): cells divided faster when glucose availability fluctuated rapidly (0.0051 division/min at a fluctuation period T = 12 min, corresponding to a doubling time of 136 minutes) than slowly (0.0027 division/min at a fluctuation period T = 192 min, corresponding to a doubling time of 257 minutes). However, this behavior was only observed in 2% glucose: the *frequency* of glucose availability did not significantly impact the division rate in 0.1% glucose (**Figure 2b**). Under periodic fluctuations of 2% glucose, the division rate was lower during half-periods without glucose than during half-periods with glucose (**Figure 2 – figure supplement 2d-f**), as expected. However, this difference depended on the frequency of glucose fluctuations: the average division rate during half-periods without glucose was higher at high frequency (small period) than at low frequency (large period) of fluctuations (**Figure 2 – figure supplement 2d-f**). Therefore, the effect of the *frequency* of glucose availability on the division rate in 2% glucose is likely due to a delay between glucose removal and growth arrest: cell proliferation never stops when the frequency of glucose depletion is too fast.

Overall, we observed two opposing patterns of cell proliferation when we varied the temporal dynamics of the metabolic environment and external osmolarity. The division rates were highest for low-frequency sorbitol fluctuations (0.0064 division/min at a fluctuation period T = 384 min) and high-frequency 2% glucose fluctuations (0.0051 division/min at a fluctuation period T = 12 min); both of these values are close to the division rate in constant 2% glucose (0.0066 division/min). Therefore, with respect to their division rate, cells behave as a low-pass filter for osmotic stress but as a high-pass filter for glucose fluctuations. Moreover, the division rate is similar when the frequencies of glucose availability and sorbitol exposure are both equal to 0.039 min^−1^ (intersection of the two curves on **Figure 2c**), corresponding to a period of 26 minutes and a division rate of 0.004 division/min. Since current models of the hyperosmotic stress response do not consider interactions with glucose metabolism, whether simultaneous fluctuations of glucose availability and osmotic stress affect cell growth additively or synergistically remains an open question. More generally, characterizing how cells respond to the dynamic phasing of two environmental components is fundamental for understanding how a living system can adapt to complex environmental changes. For these reasons, we next used our microfluidic system to quantify the division rate of cells exposed to *dual* periodic fluctuations of glucose availability and osmotic stress.

### Division rate depends on the phasing of the two stresses

To determine whether glucose availability during hyperosmotic stress impacted cell growth in dynamic conditions, we compared cell division rates under two regimes of dual periodic fluctuations that only differed in the phasing of glucose and sorbitol fluctuations. In the “in-phase stresses” (IPS) regime, glucose depletion and 1 M sorbitol stresses were applied simultaneously for half a period followed by the addition of 2% (or 0.1%) glucose and the removal of sorbitol for the other half of each period of fluctuations (**Figure 1e**). In the “alternating stresses” (AS) regime, glucose depletion and 1 M sorbitol were applied alternatively for half a period each (**Figure 1f**). We first subjected cells to dual fluctuations at a period of 24 minutes with 2% glucose, corresponding approximately to the period at which the division rate was the same when we only varied glucose availability *or* osmolarity (**Figure 2c**). Under both IPS and AS conditions, the division rate was more than 2-fold lower than under periodic fluctuations of only glucose or sorbitol (**Figure 3a**), showing that dual environmental fluctuations have a non-additive, synergistic impact on cell growth. Strikingly, cells divided about twice as fast under IPS condition (1.67 × 10^−3^ division/min, corresponding to an average doubling time of 415 minutes) than under AS condition (9.4 × 10^−4^ division/min, corresponding to an average doubling time of 737 minutes) when the fluctuation period was 24 minutes (*t*-test, P = 1.35 × 10^−5^; **Figure 3a**, **Figure 3 – figure supplement 1a,b**) or 96 minutes (2.98 × 10^−3^ division/min in IPS *vs* 1.83 × 10^−3^ division/min in AS; P = 4.10 × 10^−5^; **Figure 3b**). A similar pattern of faster growth was observed under IPS and AS conditions when we used 0.1% glucose instead of 2% glucose, for both a fluctuation period of 24 minutes (0.84 × 10^−3^ division/min under IPS *vs* 0.54 × 10^−3^ division/min under AS; *t*-test, P = 8.03 × 10^−5^; **Figure 3 – figure supplement 1c**) and 96 minutes (2.24 × 10^−3^ division/min under IPS *vs* 1.17 × 10^−3^ division/min under AS; *t*-test, P = 6.80 × 10^−3^; **Figure 3 – figure supplement 1d**). Cells also displayed strikingly different temporal dynamics of division rates under IPS and AS conditions (**Figure 3c,d**). Under IPS condition, the division rate fluctuated largely over time: after the transition to 2% glucose, the division rate quickly increased to reach a plateau (4.86 × 10^−3^ divisions/min on average during the half-period with 2% glucose); after the transition to 1 M sorbitol in the absence of glucose, cell division was greatly slowed down (1.34 × 10^−3^ divisions/minute during the half-period without glucose). In contrast, the division rate remained much more constant over time under AS condition: the average division rate was 1.69 × 10^−3^ divisions/min during the half-period with 2% glucose and 1 M sorbitol and 1.89 × 10^−3^ divisions/min during the half-period without glucose and sorbitol. Therefore, cells appear to use glucose more efficiently for growth under IPS than AS conditions. Collectively, these results further demonstrate that the timing of both glucose availability and osmotic stress matters: cells grow more slowly when facing periodic alternation of the two stresses (AS) than when facing periodic co-occurrence of these stresses (IPS).

**Caption Figure 3.**
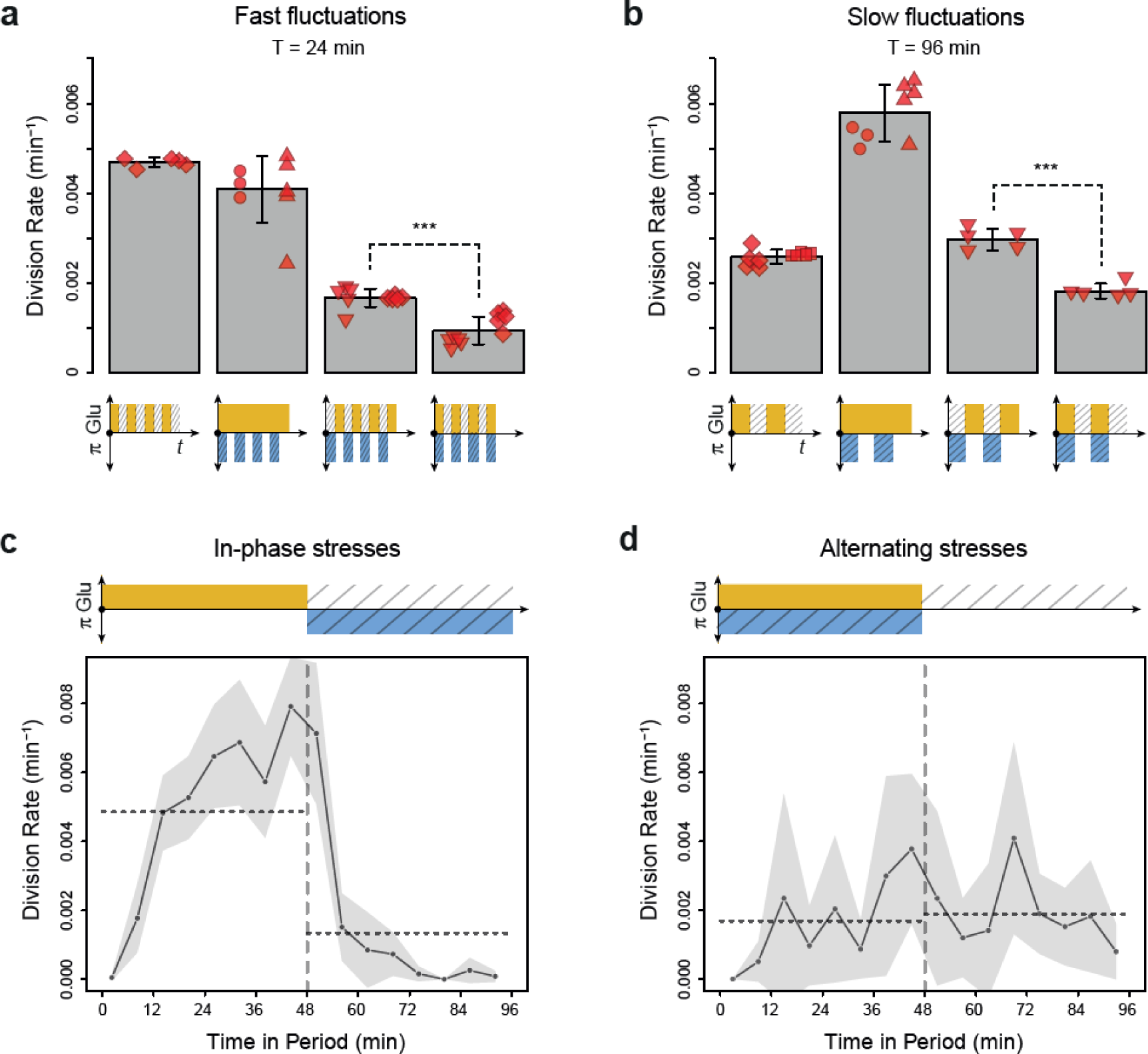
Cell division rate depends on the phasing of hyperosmotic stress and glucose availability. **(a,b)** Division rates measured in four fluctuating conditions with a period of 24 minutes and a glucose concentration of 2% (20 g/l). The four conditions are periodic glucose deprivation, periodic osmostress, in-phase stresses (IPS) and alternating stresses (AS). Bars represent mean division rates among different growth chambers. Error bars are 95% confidence intervals of the mean. Red symbols show the average division rate in each growth chamber, with different symbols representing experiments performed on different days with different microfluidic chips. Mean division rates were compared between IPS and AS conditions using *t*-tests (*** P < 0.001). **(c, d)** Temporal dynamics of division rate during a period of 96 minutes in **(c)** IPS and **(d)** AS conditions for wild-type cells. Each dot shows the average division rate during a 6-minute window centered on that dot for all fields of view sharing the same condition and for all periods in the experiment. Gray areas are 95% confidence intervals of the mean division rate. Horizontal dotted lines show the mean division rate for all data collected in each half period. The colored bars represent the periodic fluctuations in glucose (orange) and/or sorbitol (blue); hatching represents stress. **(a-d)** Cells were grown under fluctuations of 2% glucose and 1 M sorbitol. The initial number of cells analyzed among replicates ranged from 124 to 318 in **(a)** and from 94 to 342 in **(b-d)**.

The slower cell division rate observed under AS when compared to IPS could be explained by the allocation of intracellular glucose to the osmotic stress response under AS when cells are exposed to glucose and sorbitol simultaneously, leaving less glucose available for growth. Indeed, in response to hyperosmotic stress glycerol is synthesized from a glycolysis intermediate (DHAP) derived from glucose^34^. Under this hypothesis, glucose would only be fully allocated to growth in absence of hyperosmotic stress, which occurred under IPS but not AS.

### Slowdown of cell proliferation under alternating stresses is independent of HOG pathway activity

To test the hypothesis that the allocation of glucose toward glycerol synthesis explained the slower division rate observed under AS relative to IPS, we compared the division rate of mutants with impaired glycerol regulation under IPS and AS conditions. These mutant strains carried deletions of HOG1 (HOG pathway MAPK), PBS2 (MAPKK upstream of Hog1p), STE11 (MAPKKK upstream of Pbs2p), FPS1 (aquaglyceroporin regulated by Hog1p), GPD1 (glycerol-3-phosphate dehydrogenase regulated transcriptionally and post-transcriptionally by the HOG pathway) or GPD2 (paralog of GPD1). As expected, these mutants showed no growth defect in the absence of hyperosmotic stress and most mutants showed decreased division rates when exposed to constant hyperosmotic stress (**Figure 2 – figure supplement 3a-c**). At a fluctuation period of 24 minutes, the division rate was significantly lower under AS than IPS for almost all mutants (*hog1Δ*, *pbs2Δ*, *gpd1Δ*, *gpd2Δ*, *gpd1Δ; gpd2Δ* and *fps1Δ*) both with fluctuations of 2% glucose (**Figure 4a**) and 0.1% glucose (**Figure 4 - figure supplement 1a**). However, the *ste11Δ* mutant exhibited similar division rates under AS and IPS (**Figure 4a**). At a fluctuation period of 96 minutes, the division rates of the two mutants we tested, *ste11Δ* and *pbs2Δ*, were significantly lower under AS than IPS (**Figure 4 - figure supplement 1b,c**). In addition, the temporal dynamics of division rates were similar for the wild-type strain, *pbs2Δ* mutant (**Figure 4b,c**) and *ste11Δ* mutant (**Figure 4 - figure supplement 1d,e**). In conclusion, mutations known to reduce intracellular accumulation of glycerol did not attenuate the growth differences that we observed in the wild-type strain between IPS and AS conditions. Therefore, allocation of glucose toward glycerol synthesis during hyperosmotic stress is not responsible for the lower division rate observed under AS than IPS.

**Caption Figure 4.**
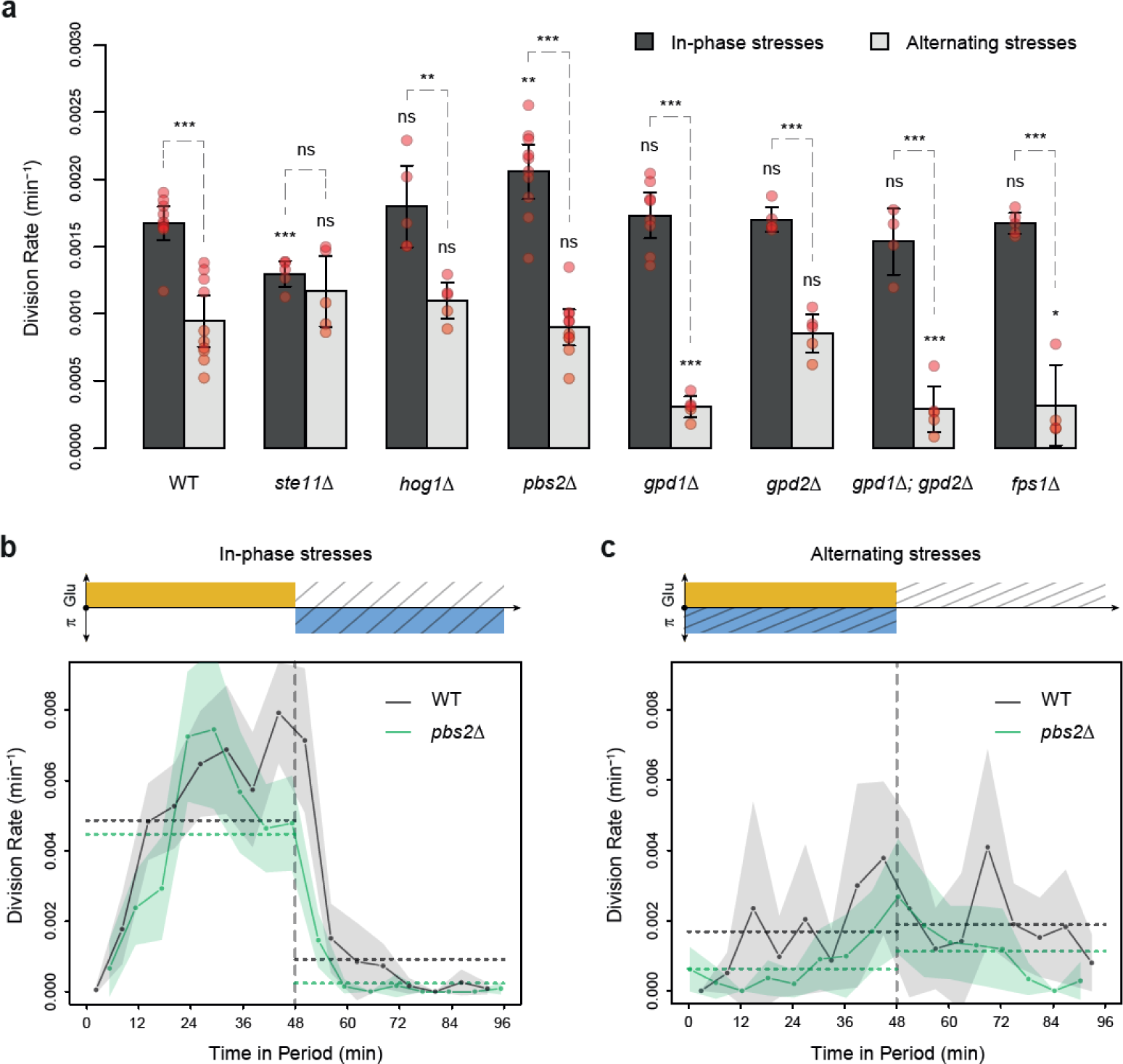
HOG pathway mutants grow faster under in-phase stresses than under alternating stresses. **(a)** Division rates measured during growth in IPS and AS conditions with 2% glucose and a fluctuation period of 24 minutes. Bars show the mean division rate measured in different growth chambers of the microfluidic chip. Error bars are 95% confidence intervals of the mean. Red symbols show the average division rate in each growth chamber. Results of *t*-tests comparing the wild-type and mutant strains under the same condition are indicated above each bar; results comparing the same strain under different conditions are shown above each pair of bars (ns: P > 0.05; * 0.01 < P < 0.05; ** 0.001 < P < 0.01; *** P < 0.001). **(b, c)** Temporal dynamics of division rate during a period of 96 minutes in **(b)** IPS and **(c)** AS conditions for wild-type (black) and *pbs2Δ* mutant (green) cells. Each dot shows the division rate during a 6-minute window centered on that dot and averaged for all fields of view sharing the same condition and all periods in the experiment. Gray and green areas are 95% confidence intervals of the mean division rate. Horizontal dotted lines show the mean division rate for all data collected in each half period. The colored bars represent the periodic fluctuations in glucose (orange) and/or sorbitol (blue); hatching represents stress. The initial number of cells analyzed among replicates ranged from 97 to 467 in **(a)** and from 124 to 145 in **(b,c)**.

### No evidence for a specific role of glucose starvation, glycogen storage or stress-induced arrest of the cell cycle in the reduced division rate observed during alternating stresses

We next tested alternative hypotheses to understand why cells grew slower under AS than IPS condition. Glucose starvation was previously shown to induce fast inhibition of transcription^35^ and translation initiation^36,37^, leading to cell growth reduction. This phenomenon may explain the slower division rate observed in AS condition than in IPS condition, because rapid arrest of the cell cycle after glucose starvation could have smaller impact on global division rate when occurring concurrently (IPS) rather than alternatively (AS) with hyperosmotic stress that also leads to fast growth reduction. We tested this hypothesis by growing wild-type cells under IPS and AS conditions with galactose instead of glucose as a carbon source, because transcriptional and translational inhibition was not observed after galactose starvation in previous studies^36,38^. We observed a significant reduction of division rate under AS condition relative to IPS condition when using galactose as a carbon source, similar to what we observed in glucose (Figure 4 – figure supplement 2). Therefore, fast inhibition of transcription and translation occurring after glucose starvation but not after galactose starvation does not contribute significantly to the slower growth in AS condition.

Yeast cells accumulate carbohydrate reserves such as glycogen to cope with nutrient starvation^39,40^, and this reserve of glycogen can be mobilized during hyperosmotic stress^41,42^. Glycogen may accumulate less under AS condition because glucose is only available during hyperosmotic stress, leading to slower growth. To test the hypothesis that glycogen storage may contribute to the difference of division rates observed between AS and IPS conditions, we quantified the division rates of *glc3Δ* mutant cells with impaired glycogen synthesis in these two conditions. Once again, we observed a significantly lower division rate of *glc3Δ* cells in AS condition relative to IPS condition similar to what was observed in wild-type cells, suggesting that glycogen storage was not significantly contributing to this difference.

Third, we tested the impact of point mutations in the cyclin inhibitor Sic1p on division rates in AS and IPS conditions. In response to hyperosmotic shock, residue 173 of Sic1p is phosphorylated by Hog1p, resulting in Sic1p stabilization and cell cycle arrest at the G1 phase^31^. Since hyperosmotic stress and glucose starvation both lead to cell growth arrest, cell division is expected to halt twice more frequently when hyperosmotic stress and glucose starvation are applied alternatively than when they are applied simultaneously, which could lead to the difference of division rates observed between AS and IPS conditions in a way that depends on Sic1p regulation. However, *sic1(T173A)* mutant cells (unphosphorylatable Sic1p) and *sic1(T172E)* mutant cells (constitutive Sic1p stabilization) showed a similar decrease of division rate in AS condition relative to IPS condition as observed in wild-type cells (**Figure 4 – figure supplement 2**). The mechanism(s) responsible for the lower division rate in AS condition relative to IPS condition therefore remains elusive.

### Cell death depends on the dynamics of the two stresses

We noticed a high proportion of wild-type cells dying under AS (**Figure 5a**): some cells suddenly burst with their nucleus staying in the growth chamber, others became opaque and stopped growing with their nucleus remaining completely still (the nucleus of living cells wobbled over time). These death events mostly occurred within minutes of the transition from medium containing 2% glucose and 1 M sorbitol to medium without glucose and sorbitol (**Figure 5 – figure supplement 1d**), suggesting cell lysis occurred due to hypo-osmotic shock following removal of 1 M sorbitol. Cell death was less frequent under IPS than AS conditions for the wild-type strain (**Figure 5a-c**), even though the frequency of hypo-osmotic shock was the same in the two conditions (**Figure 5 – figure supplement 1c,d**). We reasoned this could be due to lower intracellular accumulation of glycerol under IPS, when hyperosmotic stress is applied in the absence of glucose. Under AS, the presence of glucose during hyperosmotic stress could lead to faster intracellular accumulation of glycerol, resulting in stronger hypo-osmotic shock and cell lysis when the sorbitol concentration suddenly drops. Several pieces of evidence support this hypothesis. First, the rate of cell death should be reduced in mutants with lower glycerol synthesis. Indeed, we observed significantly lower rates of cell death for all mutants tested (*ste11Δ*, *hog1Δ*, *pbs2Δ*, *gpd1Δ*, *gpd2Δ* and *gpd1Δ; gpd2Δ*) relative to the wild-type strain under AS, but not under IPS for which glycerol synthesis is frustrated even in wild type cells due to the absence of glucose during hyperosmotic stress (**Figure 5c**). In particular, the death rate decreased from 2.4 × 10^−3^ min^−1^ for wild-type to 5.2 × 10^−5^, 3.4 × 10^−4^ and 3.5 × 10^−4^ min^−1^, respectively, for the *hog1Δ*, *pbs2Δ* and *gpd1Δ; gpd2Δ* mutants under AS when the fluctuation period was 24 minutes. A similar pattern was observed for the *pbs2Δ* mutant when the fluctuation period was 96 minutes (**Figure 5 – figure supplement 1a**), although the reduction in the death rate was less pronounced than for the period of 24 minutes. Conversely, we observed a higher death rate for the *fps1Δ* mutant (3.5 × 10^−3^ per minute) under AS (**Figure 5c**), which is consistent with higher intracellular accumulation of glycerol in this mutant lacking the Fps1 aquaglyceroporin channel involved in glycerol export. Over a 10-hour AS experiment with a fluctuation period of 24 minutes, the death rate was lowest at the beginning of the experiment and was maximal during the last five hours of the experiment (**Figure 5 – figure supplement 1e**). However, when the AS fluctuation period was 96 minutes, the maximum death rate occurred earlier (and stopped increasing after the second period) and the dynamics of cell death remained constant over multiple periods of osmotic fluctuation (**Figure 5 – figure supplement 1f**). Again, these observations are consistent with cell death being due to glycerol accumulation, since it takes time for cells to accumulate an amount of glycerol sufficient to cause bursting after hypo-osmotic shock.

**Caption Figure 5.**
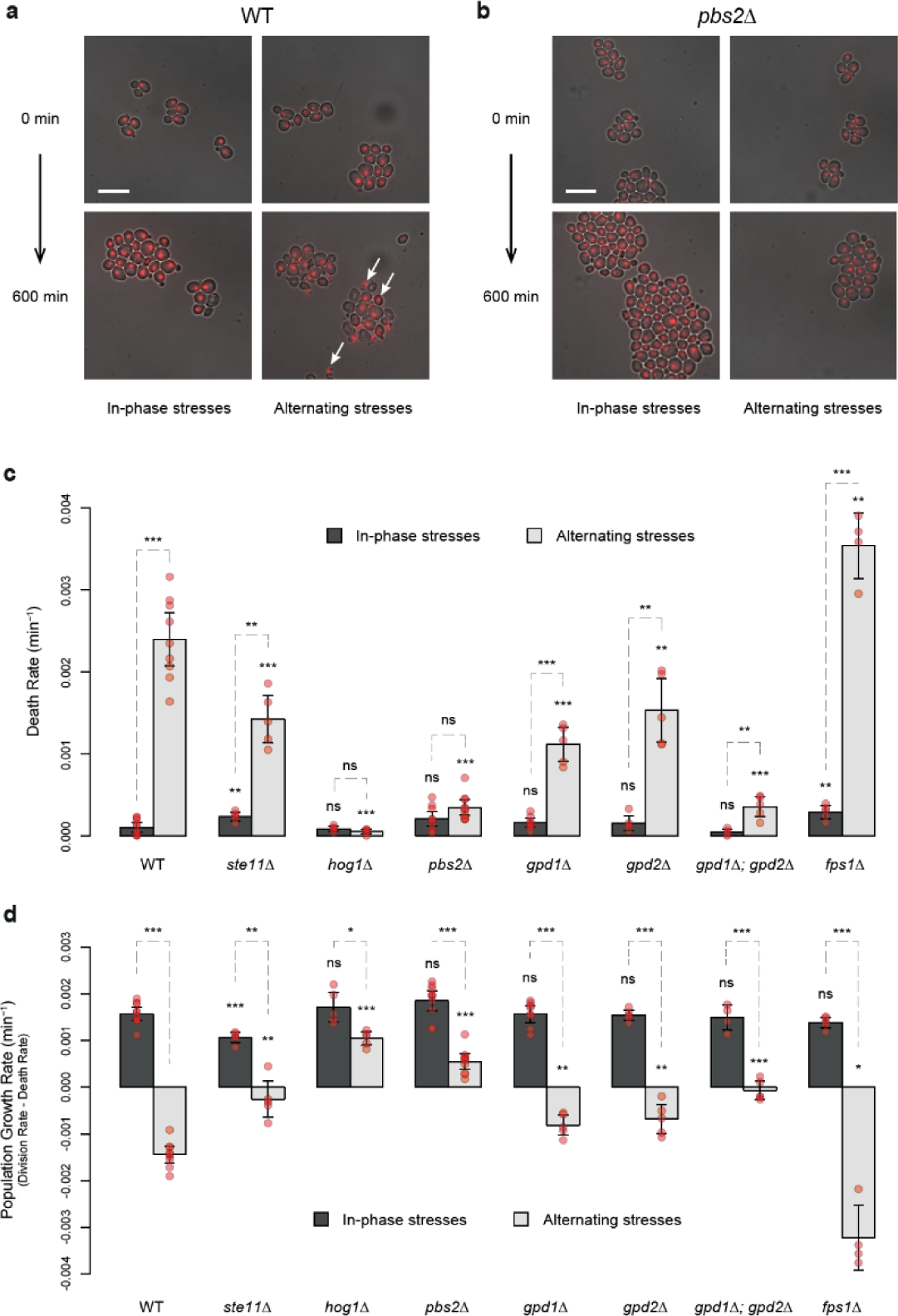
HOG pathway mutants exhibit a lower death rate and an increased population growth rate under in-phase stresses. **(a,b)** Images of wild-type **(a)** and *pbs2Δ* mutant **(b)** cells before (*t* = 0 min) and after (*t* = 600 min) growth in IPS (left) and AS (right) conditions. Fluorescence and bright field images were merged to visualize nuclei marked with HTB2-mCherry. Scale bar represents 10 µm. White arrows indicate nuclei of representative dead cells. **(c,d)** Death rates **(c)** and population growth rates **(d)** of the reference strain and seven deletion mutants under IPS and AS conditions. Population growth rates were calculated as the difference between division rates (Figure 4b) and death rates (Figure 5c). Bars show mean rates measured in different growth chambers. Error bars are 95% confidence intervals of the mean. Red symbols show the average rate for each field of view. Results of *t*-tests comparing the wild-type and mutant strains under the same conditions are indicated above each bar; results comparing the same strain under different conditions are shown above each pair of bars (ns P > 0.05; * 0.01 < P < 0.05; ** 0.001 < P < 0.01; *** P < 0.001). The initial number of cells analyzed among replicates ranged from 97 to 467 in **(c,d)**. **(a-d)** Cells were grown under fluctuations of 2% glucose and 1 M sorbitol at a period of 24 minutes.

Bonny et al.^41^ showed that *sic1* mutants (*sic1Δ* and *sic1(T173A)*) could adapt faster than wild-type cells to a hyperosmotic shock at the expense of increased cell death under repeated osmotic stresses. Consistent with their finding, we observed higher death rate of *sic1(T173A)* and *sic1(T173E)* mutant cells during repeated exposure to 1M sorbitol at a period of 24 minutes in constant 2% glucose (**Figure 5 – figure supplement 2a**). Surprisingly, we did not observe an increased death rate of these mutants under AS and IPS conditions (**Figure 5 – figure supplement 2b**), when both hyperosmotic stress and glucose availability fluctuated periodically over time. In fact, under AS condition, the death rate of *sic1(T173A)* cells was even lower than the death rate of wild-type cells. Under this condition, the particularly low division rate of *sic1(T173A)* cells may lead to strengthened cell wall, decreasing the probability of cell bursting after hypo-osmotic shocks.

### HOG pathway mutants are fitter than wild-type cells under fast alternating stresses

The rates of cell division and cell death both contribute to fitness (*i.e.,* the adaptive value) of a genotype in a particular environment. Since HOG pathway mutants exhibited different cell division and death rates compared to the wild-type genotype under AS, we calculated the population growth rate (division rate minus death rate) as a fitness estimate. Under IPS, the population growth rates of most mutant strains and of the wild-type strain were not significantly different; the only exception being the slightly lower growth rate of the *ste11Δ* mutant (**Figure 5d**). However, under AS with a fluctuation period of 24 minutes, several mutants had higher population growth rates than the wild-type strain (**Figure 5d**). In fact, the population growth rate was negative for the wild-type strain (−1.4 × 10^−3^ min^−1^) as cells died faster than they divided and positive for the *hog1Δ* mutant (1.0 × 10^−3^ min^−1^) and *pbs2Δ* mutant (5.5 × 10^−4^ min^−1^). These differences are clear in the microscopy images, as the population of wild-type cells visually shrank over time under AS (**Figure 5a**), while the population of *pbs2Δ* cells clearly expanded (**Figure 5b**). Thus, the *hog1Δ* and *pbs2Δ* genotypes are better adapted and would quickly outcompete wild-type cells under these dynamic conditions. However, this is only true when the frequency of environmental fluctuations is sufficiently high, since we did not observe significant differences in the population growth rate between the wild-type and *pbs2Δ* mutant under AS when the fluctuation period was 96 minutes (**Figure 5 – figure supplement 1b**). We conclude that mutants that were first characterized by an inability to adapt to prolonged hyperosmotic stress can be well adapted when hyperosmotic stress rapidly fluctuates in antiphase with glucose availability. Therefore, the genetic mechanisms that contribute to adaptation under steady-state conditions could be detrimental under dynamic conditions, highlighting the importance of investigating how organisms adapt to dynamically changing environments.

### Osmoregulation is impaired under in-phase stresses but not under alternating stresses

The ability of cells to sense environmental fluctuations and to execute an adaptive response has been mainly studied using fluctuations of one stress cue at a time. How cells sense and respond to dual fluctuations of two interacting stresses remains a fundamental open question to understand how cells cope with complex environmental dynamics. Our findings suggest that the cell response to dual stress fluctuations can be very different depending on the phasing of the two stresses. Indeed, yeast cells appear to accumulate more glycerol under alternating stresses (AS) than under in-phase stresses (IPS). This could be either because of an impaired ability of cells to sense hyperosmotic shocks in absence of glucose or because of an impaired capacity to respond to hyperosmotic shocks in absence of glucose. Glycerol synthesis is regulated by the HOG pathway; thus, we investigated whether the activity of this pathway differed under IPS and AS conditions.

In the presence of glucose, activation of the HOG pathway in response to hyperosmotic shock triggers phosphorylation and nuclear translocation of Hog1p, which regulates transcription of multiple genes. A negative feedback loop dephosphorylates Hog1p, which leaves the nucleus in less than 15 minutes even if hyperosmotic stress is maintained^4,21^. To track HOG pathway activity, we quantified the nuclear enrichment of Hog1p over time by monitoring the subcellular location of Hog1 protein fused to a fluorescent marker (Hog1-GFP) in cells that also expressed the nuclear marker Htb2-mCherry. We did not detect any enrichment of Hog1-GFP in nuclei when only glucose fluctuated over time (**Figure 6 – figure supplement 1a**), suggesting cells did not perceive glucose fluctuations as a significant osmotic cue. Although previous studies observed small transient (less than two minutes) peaks of Hog1-GFP nuclear localization after glucose was added back to the medium following glucose depletion^23,43^, the temporal resolution in our experiments (one image every 6 minutes) may have been too low to detect these peaks.

**Caption Figure 6.**
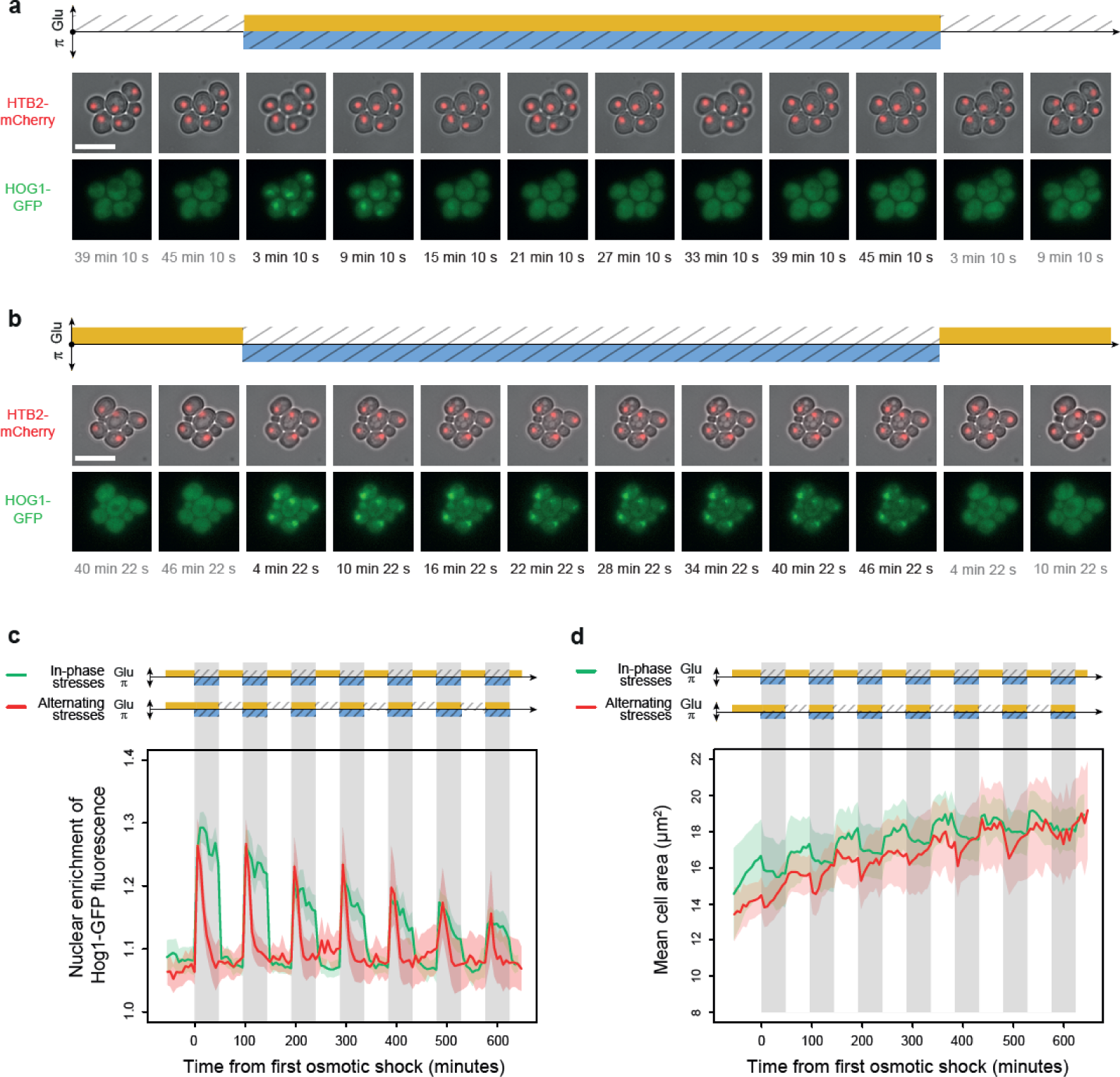
Osmoregulation is delayed after hyperosmotic stress under in-phase stresses but not under alternating stresses. **(a-b)** Timelapse images of cells expressing Htb2-mCherry and Hog1-GFP under AS **(a)** and IPS conditions **(b)** for periods of 96 minutes, showing cellular localization of Hog1p during the second osmotic shock in each experiment. Top: fluorescence and brightfield images merged to visualize cell nuclei tagged with histone HTB2-mCherry. Bottom: fluorescence images showing Hog1-GFP localization. The time since the last environmental change is indicated below each image. Scale bars represent 10 µm. **(c)** Temporal dynamics of the enrichment of Hog1-GFP fluorescence in cell nuclei under IPS (red curve) and AS (green curve) conditions. Colored areas indicate 95% confidence intervals. **(d)** Temporal dynamics of cell size (area) in IPS (green) and AS (red) conditions for the same cells as in panel (**c)**. Each curve shows the mean area measured among cells. Colored areas indicate 95% confidence intervals of the mean. **(c-d)** Each curve shows the mean nuclear enrichment or mean cell size for 11 to 25 cells in one or two fields of view. The colored graphs represent the periodic fluctuations of glucose (orange) and/or sorbitol (blue); hatching represents stress; gray indicates exposure to 1 M sorbitol. **(a-d)** Cells were grown under fluctuations of 2% glucose and 1 M sorbitol at a period of 96 minutes.

In contrast, enrichment of Hog1-GFP fluorescence in nuclei was observed within minutes after exposure to 1 M sorbitol under both AS (**Figure 6a**) and IPS (**Figure 6b**). Therefore, cells can sense hyperosmotic shock, activate the HOG MAPK cascade and phosphorylate Hog1 MAP kinase both in the presence (AS) and absence (IPS) of glucose. However, the adaptation dynamics of Hog1p (*i.e.,* its exit from the nucleus) were remarkably different (**Figure 6a-c**): under AS, nuclear enrichment of Hog1-GFP peaked at 6 minutes following hyperosmotic shock and then quickly decayed and became undetectable after 30 minutes (**Figure 6a,c**)—essentially the same dynamics observed under periodic fluctuations of osmotic stress without glucose fluctuations ((**Figure 6 – figure supplement 1a**). Under IPS conditions, nuclear enrichment also peaked 6 minutes after hyperosmotic shock, but Hog1-GFP returned to the cytosol much more slowly; strong nuclear enrichment was still observed 48 minutes after exposure to hyperosmotic stress in the absence of glucose (**Figure 6b,c**). When the hyperosmotic stress was released, Hog1-GFP returned to the cytosol in less than 12 minutes.

To determine whether Hog1-GFP eventually returns to the cytoplasm during hyperosmotic stress in the absence of glucose, we applied a single pulse of 1 M sorbitol without glucose for 4 hours. Nuclear enrichment of Hog1-GFP reached basal levels about two hours after the onset of hyperosmotic stress (**Figure 6 – figure supplement 1b**). The delayed exit of Hog1-GFP out of the nucleus under IPS suggests the activity of the feedback loop regulating Hog1p dephosphorylation is impaired in the absence of glucose. We hypothesized that delayed nuclear export of Hog1-GFP under IPS could be due to impaired osmoregulation. In support of this hypothesis, we observed no recovery of cell size during hyperosmotic stress under IPS conditions (**Figure 6d**). Cells only returned to their initial size when sorbitol was removed and glucose was added back to the medium, which also corresponded to the moment when Hog1-GFP returned to the cytosol. In contrast, under AS, both the recovery of cell size and nuclear export of Hog1-GFP occurred while cells were still exposed to hyperosmotic stress (**Figure 6**), showing that osmoregulation was not impaired under these conditions. These results suggest glucose is necessary for the rapid osmoregulation that usually occurs in the first 20 minutes following hyperosmotic stress. This fast osmoregulation has been proposed to rely on the induction of glycerol synthesis via Hog1p-dependent post-translational mechanisms^26^. Since glucose is a metabolic precursor of glycerol, the absence of glucose may prevent glycerol synthesis and thereby fast osmoregulation. Further work will be necessary to test this hypothesis and study how glucose stored in the cell is used (or not) for glycerol production.

### Transcriptional response is impacted by the interaction between two environmental dynamics

Temporal fluctuations of two different stresses may have different or even opposite effects on gene expression, raising the question of how dual fluctuations of these two stresses would affect gene expression. Fast periodic fluctuation of osmotic stress was previously shown to cause hyper-activation of the STL1 promoter (P_STL1_) regulated by Hog1p^27^, while glucose depletion is known to inhibit transcription^35^ and translation initiation^44,45^. We therefore asked whether dual fluctuations of glucose depletion and osmotic stress had additive effects on P_STL1_ expression or whether one of the dynamic cues had a dominant effect. To address this question, we quantified the expression dynamics of a *P_STL1_-mCitrine* fluorescent reporter gene under IPS and AS conditions. As expected, we observed transient expression of *P_STL1_-mCitrine* in response to both short (48 min) and long (10 h) pulses of 1 M sorbitol (**Figure 7**): the mean fluorescence level peaked at a 2.8-fold change 115 minutes after exposure to a short sorbitol pulse and 2.4-fold change 122 minutes after a long sorbitol pulse; then, fluorescence gradually returned to basal levels, even when osmotic stress was maintained. This expression pattern is consistent with previous quantifications of P_STL1_ transcriptional activity during hyperosmotic stress^46,47^. When hyperosmotic shocks were periodically applied for 48 minutes every 96 minutes, *P_STL1_-mCitrine* expression constantly increased to reach 10.2-fold change after 11 hours (**Figure 7b**), suggesting that osmoadaptation never occurred under this condition. Interestingly, hyper-activation of *P_STL1_-mCitrine* transcription was also observed under AS conditions where *P_STL1_-mCitrine* expression reached a maximum of 4.1-fold change after 413 minutes and remained as high as 3.1-fold-change after 11 hours under the AS regime with a fluctuation period of 96 minutes (**Figure 7a,b**).

**Caption Figure 7.**
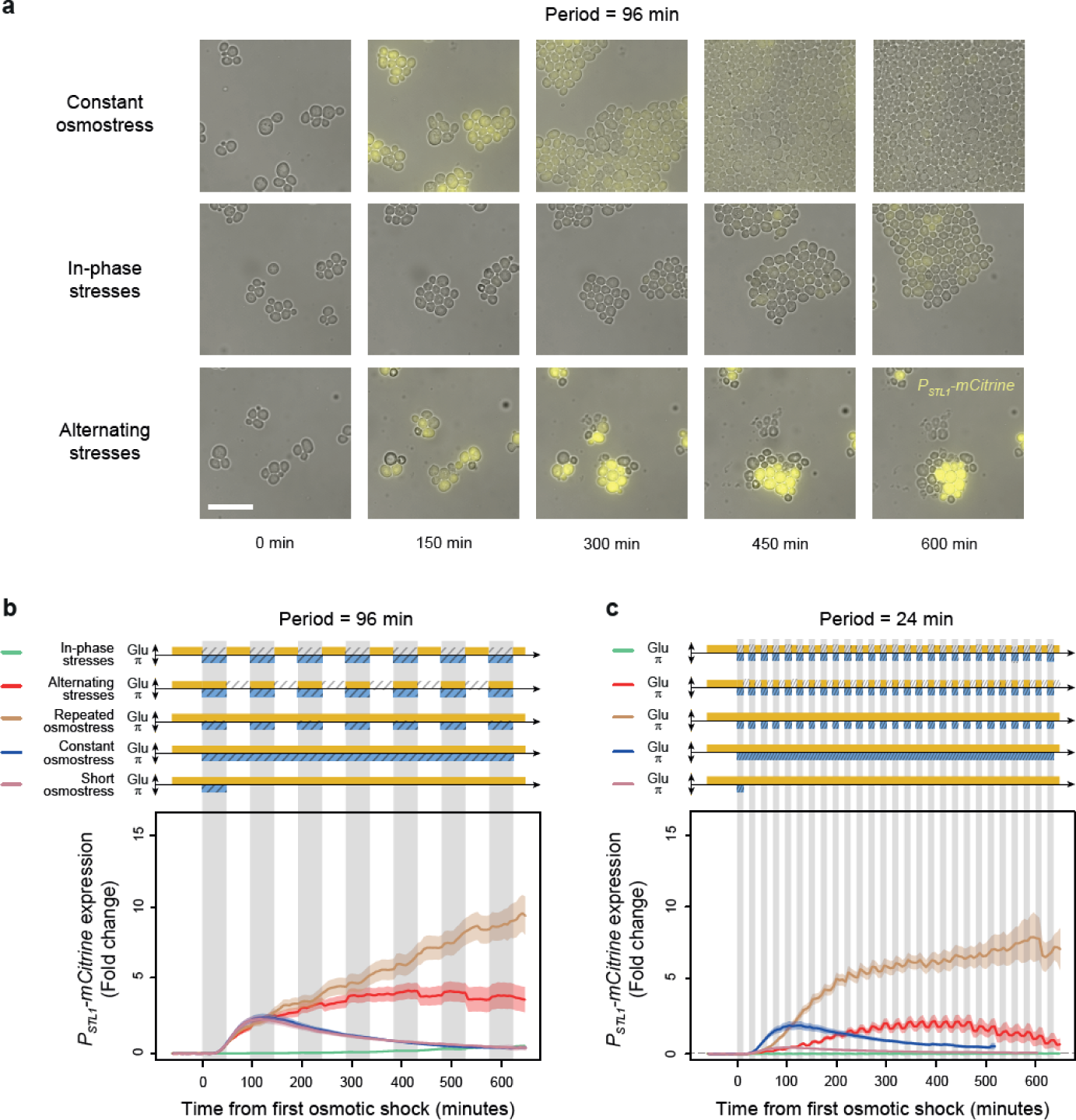
A transcriptional target of the HOG pathway is under-expressed during in-phase stresses and over-expressed during alternating stresses. **(a)** Images of cells expressing a fluorescent reporter (mCitrine) under control of the STL1 promoter known to be regulated by the HOG pathway. Rows correspond to different conditions (as indicated on the left) and columns correspond to different time points after transitioning from complete medium to each condition. Bright field and fluorescence images are overlaid. The scale bar represents 20 µm. **(b-c)** Temporal dynamics of *P_STL1_-mCitrine* expression in five conditions: IPS (green curves), AS (red curve), periodic osmotic stress with constant glucose (brown curve), a single transition to constant osmotic stress with glucose for 10 hours (blue curve), and a short pulse of osmotic stress with constant glucose (purple curve). The fluctuation period is 96 minutes in **(b)** and 24 minutes in **(c)** for the first three conditions. 2% glucose was used in all conditions. Each curve shows the mean fold change of fluorescence intensity measured for 30—65 cells from four or five fields of view. Colored areas indicate 95% confidence intervals of the mean. The colored graphs represent the periodic fluctuations of glucose (orange) and/or sorbitol (blue); hatching represents stress; gray indicates exposure to 1 M sorbitol.

Conversely, we observed much weaker and slower induction of *P_STL1_-mCitrine* expression under IPS: the maximal fold change was 0.7 after 11 hours of IPS with a fluctuation period of 96 minutes (**Figure 7a,b**). We observed a similar pattern (*i.e.,* faster, stronger induction of *P_STL1_-mCitrine* under AS than IPS) when the fluctuation period was 24 minutes (**Figure 7c**). Therefore, the STL1 promoter is not activated by hyperosmotic stress in the absence of glucose, despite nuclear translocation of the MAP kinase Hog1p. This result suggests that the global repression of expression in response to abrupt glucose starvation is dominant over the hyper-activation of P_STL1_ transcriptional activity induced by periodic osmotic stress.

## Discussion

Using microfluidics and time lapse microscopy, we studied how yeast cells behave when confronted with two dynamic stresses that were applied either simultaneously (in-phase) or alternatively (in antiphase). This work demonstrates the value of applying artificial periodic fluctuations in several environmental parameters to understand how a biological system can process and integrate information from multiple cues in its environment. By using periodic inputs, we were able to investigate how changing the phasing of two stresses impacted fitness while keeping the duration of each stress constant. We found that dual fluctuations of glucose deprivation and osmotic stress had synergistic (non-additive) effects on cell proliferation that could not be easily predicted from the effects of fluctuating each stress separately. The phasing of the two stresses had a striking impact on several cell phenotypes, including division rate, death rate, osmoregulation and transcriptional activation of a gene regulated by osmotic stress. The quantitative measurements of fitness that we collected for diverse genotypes under dynamic conditions with different glucose (main carbon source) concentrations can be used to further constrain mathematical models of yeast stress responses by adding two important features: resource allocation and regulation based on the presence of glucose. Moreover, our results indicate that the classic picture of yeast adaptation to osmotic stress cannot fully explain the behavior of yeast cells under fluctuating conditions. Indeed, by making glucose available for only short durations, we produced an environment in which the dynamics of the osmotic signaling pathway and its interaction with the glucose sensing pathway and glycolysis are critical. While the HOG signaling pathway was activated under all conditions, transcription of the target genes and inactivation of the HOG pathway were only detected when glucose and hyperosmotic stress were applied simultaneously, but not when hyperosmotic stress occurred in the absence of glucose. Therefore, if glucose is not available in the environment, cells are unable to commit to classical transcriptional/translational responses after osmotic stress. This finding may seem inconsistent with conclusions of a recent study showing that yeast cells displayed a stronger response to hyperosmotic stress at lower glucose concentration, including higher expression of several genes regulated by the Hog pathway^48^. However, key differences in experimental procedures may explain this apparent discrepancy: Shen et al.^48^ investigated the response to a single hyperosmotic shock at diverse constant glucose concentrations – not dynamic stresses as we did – and the lowest glucose concentration they considered was 0.02% allowing slow cell growth - not complete glucose depletion that stopped growth. This further suggests that subtle changes in the metabolic environment of cells may have profound impact on their stress-response capacity.

Our findings shed lights on the importance of including different types of metabolic environment fluctuations when describing stress responses and probing gene regulatory networks with time varying signals to constrain both mathematical and biological models of stress responses. Interestingly, another recent study showed that the interplay between osmotic stress and glucose concentration could cause bimodal expression of a key determinant of cell survival after starvation^49^. Since hyperosmotic stress was induced using high glucose concentration in this previous study, it would be particularly interesting to determine whether periodic fluctuations of hyperosmotic glucose concentration caused similar fitness defects as we observed when periodic fluctuations of glucose and sorbitol were applied in phase.

Importantly, in contrast to the well-known proliferative defects of osmo-sensitive mutants under constant hyperosmotic conditions, we show that periodically varying osmotic stress is not always detrimental to the growth of osmo-sensitive yeast cells. Indeed, HOG pathway mutants can grow and are even fitter than wild-type cells under fast alternating fluctuations of glucose deprivation and hyperosmotic stress. In this condition, osmo-sensitive mutant cells survive better than wild-type cells to repeated hypo-osmotic shocks, probably because they do not accumulate glycerol in response to hyperosmotic shocks and thus are less sensitive to fluctuations in the osmolarity of their environment. This also suggests cells can be killed by exploiting their adaptative response. Forcing cells to repeatedly express their osmoadaptation program makes them very sensitive to osmotic rupture of the cell wall. Hence, periodically stressing cells (and their metabolic state) may be an efficient strategy to kill yeast cells. We further imagine that varying the timescale of fluctuations may even prevent cells from finding an evolutionary escape route.

Overall, we anticipate the importance of extending our study of the interaction of metabolic resources with other critical stresses, including antibiotic or mechanical stress, to higher eukaryotes. We propose that sending periodically a metabolic resource combined or alternating with a stressing agent allows to study the interplay between stress response and cell metabolism. We anticipate that such study could open novel research areas by revisiting the questions of stress response dynamics, within a systems view of cells homeostasis, in which metabolic activity regulates or interferes with multiple important cellular processes.

## Materials and Methods Summary

All *Saccharomyces cerevisiae* strains used in this study are derived from BY4741 or BY4742^50^ and are listed in Supplementary Tables 1 and 2. Strains made in this study were obtained primarily using CRISPR/Cas9 gene editing. Yeast cells were cultivated and imaged in custom-made microfluidic devices for all time-lapse microscopy experiments described in this study. The microfluidic chip consists of five independent pairs of flow channels (800 µm wide x 50 µm high) connected each to five growth chambers (400 x 400 x 3.8 µm; L x W x H) where yeast cells are constrained to proliferate in monolayer. A custom-made valve controller was used to dynamically control which medium was dispensed to the cells based on a predetermined schedule of valve state switching. The chip was then mounted on a motorized inverted microscope (Olympus IX83) equipped with LEDs for fluorescence excitation (CoolLED pE-300ultra), a Zyla 4.2 sCMOS camera (Andor - Oxford Instruments) and an autofocus module (IX3-ZDC2, Olympus). The microscope was controlled using iQ v3.6.3 software (Andor Technology) and all images were obtained using a 60x oil immersion objective (Olympus PlanApo N 60X/1.42) and a 1.6x magnification changer. Additional details on cell construction, cell culture, image acquisition and image analysis can be found in the supplementary materials and methods.

## Acknowledgments

The authors would like to thank their team members for their critical reading of this manuscript. We also thank Williams Brett who helped us design the microfluidic control system. This work was supported by the European Research Council grant SmartCells (724813) and received support from grants ANR-11-LABX-0038, ANR-10-IDEX-0001-02 and ANR-16-CE12-0025-01.

## Author Contributions

FD, CC, AL performed experiments; FD, CC, AL, MLB, JS performed strain construction; FD, BS, SP, MLB, CC, LC, JMdM, PH analyzed the data and wrote the manuscript; FD and PH designed the project.

## Supplementary Information

Supplementary information contains 7 supplementary figures and detailed material and methods.

## Supplementary Files

**Supplementary File 1.** R scripts used to perform all analyses and to generate all figures.

**Supplementary File 2.** Source data for all figures. Files named “Data.xxx” contain values for all samples included in each figure. Files named “Summary.Data.xxx” contain values averaged among replicates and standard deviation of replicates.

Supplementary files are also available on the following Zenodo public repository 10.5281/zenodo.10471016

