## supplementary file for "Yeast cell responses and survival during periodic osmotic stress are controlled by glucose availability"

### Materials and Methods

#### Yeast strains

All *Saccharomyces cerevisiae* strains used in this study are derived from BY4741 or BY4742[1] and are listed in Supplementary Tables 1 and 2 below. The reference strain yPH_132 expresses the mCherry fluorescent reporter fused to histone H2B to label cell nuclei. To obtain this strain, the mCherry_pAgTEF-KanMX4-tAgTEF linear DNA fragment was amplified from plasmid pYM35 (PCR_TOOLBOX collection from EUROSCARF). This DNA fragment was inserted at the *HTB2* locus in strain BY4741 using a classic LiAc transformation protocol[2] and a clone resistant to G418 was stored at -80°C as strain yPH_132.

All gene-deletion strains of yeast used in this study were obtained by directed mutagenesis of the yPH_132 strain (*HTB2::HTB2-mCherry*). Gene deletions were performed using CRISPR/Cas9 gene editing according to the method described in Laughery et al.[3] that relies on the co-transformation of i) a plasmid vector derived from pML104 (Addgene #67638) allowing expression of the Cas9 protein and a guide RNA specific to the target gene and ii) a DNA repair fragment designed to delete the coding sequence of the target gene via homology directed repair. For each gene deletion, a sequence containing the 20 nucleotides of the sgRNA recognition site in the coding sequence of the target gene was cloned between BclI and SwaI restriction sites in the pML104 plasmid by ligation of hybridized oligonucleotides. The DNA repair fragment was obtained by PCR amplification (Phusion® Hot Start Flex 2X Master Mix, NEB M0536S) of two oligonucleotides with 20 bases of reverse complementarity to each other at the 3′ end and 70 bases homologous to the region either immediately upstream or downstream of the coding sequence of the target gene at the 5′ end. The repair fragment and the CRISPR/Cas9 plasmid specific to each gene were transformed together in exponentially growing yPH_132 cells using a standard lithium acetate method. Transformants were isolated on CSM-ura (Formedium™, DCS0271) agar plates and gene deletions were confirmed by PCR screening and by Sanger sequencing. Positive clones were grown on YPG plates (10 g/L Yeast extract, 20g/L peptone, 5% (v/v) glycerol, 20 g/L bacto agar) to counter select petite cells (*ρ*- phenotype) and then transferred on CSM agar plates containing 0,8 g/L 5-Fluoroorotic Acid (Thermo Scientific™ R0812) to counter select the CRISPR/Cas9 plasmid carrying the Ura3 gene. Two independent clones were stored at -80°C in 15% glycerol for each gene deletion (the clones used in this study are listed in Supplementary Table 1).

**Supplementary Table 1.** Genotype of yeast strains used in this study.

| Name | Genotype | Mating type | Background | Reference or source |
|---|---|---|---|---|
| yPH_132 | *HTB2::HTB2-mCherry_pTEF-KanMX4-tTEF his3Δ1 leu2Δ0 met15Δ0 ura3Δ0* | a | BY4741 | This work |
| yPH_015 | *HTB2::mCherry-Ura3 HOG1-GFP* | a | BY4741 | Our team |
| yPH_091 | *pSTL1::yECITRINE-HIS5 his3Δ1 leu2Δ0 lys2Δ0 ura3Δ0 Hog1::mCherry-hph* | alpha | BY4742 | https://doi.org/10.1073/pnas.1206810109 |
| yPH_051 | *pGPD1::YFP* | a | BY4741 | Gift from Megan McClean |
| yPH_403 | *HTB2::HTB2-mCherry_pTEF-KanMX4-tTEF pbs2Δ his3Δ1 leu2Δ0 met15Δ0 ura3Δ0* | a | yPH_132 | This work |
| yPH_405 | *HTB2::HTB2-mCherry_pTEF-KanMX4-tTEF fps1Δ his3Δ1 leu2Δ0 met15Δ0 ura3Δ0* | a | yPH_132 | This work |
| yPH_412 | *HTB2::HTB2-mCherry_pTEF-KanMX4-tTEF gpd1Δ his3Δ1 leu2Δ0 met15Δ0 ura3Δ0* | a | yPH_132 | This work |
| yPH_414 | *HTB2::HTB2-mCherry_pTEF-KanMX4-tTEF gpd2Δ his3Δ1 leu2Δ0 met15Δ0 ura3Δ0* | a | yPH_132 | This work |
| yPH_421 | *HTB2::HTB2-mCherry_pTEF-KanMX4-tTEF gpd1Δ gpd2Δ his3Δ1 leu2Δ0 met15Δ0 ura3Δ0* | a | yPH_132 | This work |
| yPH_441 | *HTB2::HTB2-mCherry_pTEF-KanMX4-tTEF hog1Δ his3Δ1 leu2Δ0 met15Δ0 ura3Δ0* | a | yPH_132 | This work |
| yPH_445 | *HTB2::HTB2-mCherry_pTEF-KanMX4-tTEF ste11Δ his3Δ1 leu2Δ0 met15Δ0 ura3Δ0* | a | yPH_132 | This work |
| yPH_452 | *HTB2::HTB2-mCherry_pTEF-KanMX4-tTEF glc3Δ his3Δ1 leu2Δ0 met15Δ0 ura3Δ0* | a | yPH_132 | This work |
| yPH_482 | *HTB2::HTB2-mCherry_pTEF-KanMX4-tTEF sic1(T173A) his3Δ1 leu2Δ0 met15Δ0 ura3Δ0* | a | yPH_132 | This work |
| yPH_486 | *HTB2::HTB2-mCherry_pTEF-KanMX4-tTEF sic1(T173E) his3Δ1 leu2Δ0 met15Δ0 ura3Δ0* | a | yPH_132 | This work |

**Supplementary Table 2.** Oligonucleotides used to generate gene deletions and point mutations.

| Strain | Primer name | Sequence |
|---|---|---|
| *pbs2Δ* (yPH_403) | sgRNA_PBS2_F | GATCAATCAAAGCGAGCAAGACAAGTTTTAGAGCTAG |
| | sgRNA_PBS2_R | CTAGCTCTAAAACTTGTCTTGCTCGCTTTGATT |
| | Deletion_PBS2_F | CGTCATACAACTAAAACTGATAAAGTACCCGTTTTTCCGTACATTTCTATAGATACATTATTATATTAAGCAGATCGAGACGTTAATTTC |
| | Deletion_PBS2_R | GTAGCTTTTCGTCTGCTTTTTTTTTGTTGTTATATTCACGTGCCTGTTTGCTTTTATTTGGATATTAACGGAAATTAACGTCTCGATCTG |
| | Seq_PBS2_F | GCTTACCTGCTTGCCGGAAG |
| | Seq_PBS2_R | CTATAACGAGTATAATGCAAG |
| *gpd1Δ* (yPH_412) (yPH_421) | sgRNA_GPD1_F | GATCTCTGCTGCCATCCAAAGAGTGTTTTAGAGCTAG |
| | sgRNA_GPD1_R | CTAGCTCTAAAACACTCTTTGGATGGCAGCAGA |
| | Deletion_GPD1_F | TATACTACCATGAGTGAAACTGTTACGTTACCTTAAATTCTTTCTCCCTTTAATTTTCTTTTATCTTACTCTCCTACATAAGACATCAAG |
| | Deletion_GPD1_R | ATGAATATGATATAGAAGAGCCTCGAAAAAAGTGGGGGAAAGTATGATATGTTATCTTTCTCCAATAAATCTTGATGTCTTATGTAGGAG |
| | Seq_GPD1_F | GCACAACAAGTATCAGAATG |
| | Seq_GPD1_R | ATGCGGAAGAGGTGTACAGC |
| *gpd2Δ* (yPH_414) (yPH_421) | sgRNA_GPD2_F | GATCGCATTGGTCCGAAACCACCGGTTTTAGAGCTAG |
| | sgRNA_GPD2_R | CTAGCTCTAAAACCGGTGGTTTCGGACCAATGC |
| | Deletion_GPD2_F | TTTTTTTTTATATATTAATTTTTAAGTTTATGTATTTTGGTAGATTCAATTCTCTTTCCCTTTCCTTTTCCTTCGCTCCCCTTCCTTATC |
| | Deletion_GPD2_R | ATAATGATAAATTGGTTGGGGGAAAAAGAGGCAACAGGAAAGATCAGAGGGGGAGGGGGGGGGAGAGTGTGATAAGGAAGGGGAGCGAAG |
| | Seq_GPD2_F | CAGCTCTTCTCTACCCTGTC |
| | Seq_GPD2_R | GGTGATGTGATATGTAAACG |
| *fps1Δ* (yPH_405) | sgRNA_FPS1_F | GATCACAGCAGGACAATTTCAACGGTTTTAGAGCTAG |
| | sgRNA_FPS1_R | CTAGCTCTAAAACCGTTGAAATTGTCCTGCTGT |
| | Deletion_FPS1_F | ATCAACAAAGTATAACGCCTATTGTCCCAATAAGCGTCGGTTGTTCTTCTTTATTATTTTACCAAGTACGCTCGAGGGTACATTCTAATG |
| | Deletion_FPS1_R | TACCGGCGGTAGTAAGCAGTATTTTTTTCTATCAGTCTATATTATTTGTTTCTTTTTCTTGTCTGTTTTCCATTAGAATGTACCCTCGAG |
| | Seq_FPS1_F | CAGTGTGAATCCGGAGACGG |
| | Seq_FPS1_R | TACTTAAGACGATGGGTCAG |
| *hog1Δ* (yPH_441) | sgRNA_HOG1_F | GATCGGCTCCTTACCACGATCCAAGTTTTAGAGCTAG |
| | sgRNA_HOG1_R | CTAGCTCTAAAACTTGGATCGTGGTAAGGAGCC |
| | Deletion_HOG1_F | TGGTAAATACTAGACTCGAAAAAAAGGAACAAAGGGAAAACAGGGAAAACTACAACTATCGTATATAATAGTCCCTAACCACTCATTCTT |

| | | |
|---|---|---|
| | Deletion_HOG1_R | TTCCTCTATACAACTATATACGTAAATACTTTTATGAGTACCATAAAAAAAAGAAACATCAAAAAGAAGTAAGAATGAGTGGTTAGGGAC |
| | Seq_HOG1_F | TAGTGGAAGAGGAATTTGCG |
| | Seq_HOG1_R | GCCATAAGTGACGGTTCTTG |
| *ste11Δ* (yPH_445) | sgRNA_STE11_F | GATCTATGGTGCTTCTCAAGAAGGGTTTTAGAGCTAG |
| | sgRNA_STE11_R | CTAGCTCTAAAACCCTTCTTGAGAAGCACCATA |
| | Deletion_STE11_F | CAGTAGAAAATATTCATATTTACACACATGCATAAAGAGAGACCACTTAATAAAGCTAGTATGATAAGATCACCGGTAGACGAAATATAC |
| | Deletion_STE11_R | ATGTATTATTTGATAAAAAATCGGCCAGAGCACTTTAGTGCCATAAAAAGAATTAATAAGTAGCCCTTTTGTATATTTCGTCTACCGGTG |
| | Seq_STE11_F | TTCTTTATGCTGTCCTCACC |
| | Seq_STE11_R | GAGAATCAAATACCGTCATC |
| *glc3Δ* (yPH_452) | sgRNA_GLC3_F | GATCTTTCGACTACAGATTAGCAAGTTTTAGAGCTAG |
| | sgRNA_GLC3_R | CTAGCTCTAAAACTTGCTAATCTGTAGTCGAAA |
| | Deletion_GLC3_F | TCCTACATTTTTTTTCCCTGATAACTTCCTGTTACTATTTAAGAACACCAAACCAAGTATAAAGAACCGTCAAGAATAAAACTCTATACT |
| | Deltetion_GLC3_R | GTACGTTTAGATATCTACCAATACATGAAGAGAAAAAAATTATTGAGTCTTGATTTTCAGTAAGCAATATAGTATAGAGTTTTATTCTTG |
| | Seq_GLC3_F | TCGAGCCAAGTGACACCAGC |
| | Seq_GLC3_R | GACAGCTCTGCTATTCGCCC |
| *Sic1* (yPH_482 T173A) (yPH_486 T173E) | sgRNA_SIC1_F | GATCACCTGGTACGCCCAGCGACAGTTTTAGAGCTAG |
| | sgRNA_SIC1_R | CTAGCTCTAAAACTGTCGCTGGGCGTACCAGGT |
| | Repair_SIC1_T173A_F | ACATTTATCACTTGAAAGAGATGAGTTTGATCAGACACATAGAAAGAAGATTATTAAAGATGTACCTGGTGCGCCCAGCGACAAAGTGAT |
| | Repair_SIC1_T173A_R | TTCACTTTCTTGACTCCTGGCGTCATTTTTCGGAGAGTTGTTGTTCCAATTTTTTGCCAATTCAAATGTTATCACTTTGTCGCTGGGCGC |
| | Repair_SIC1_T173E_F | ACATTTATCACTTGAAAGAGATGAGTTTGATCAGACACATAGAAAGAAGATTATTAAAGATGTACCTGGTGAGCCCAGCGACAAAGTGAT |
| | Repair_SIC1_T173E_R | TTCACTTTCTTGACTCCTGGCGTCATTTTTCGGAGAGTTGTTGTTCCAATTTTTTGCCAATTCAAATGTTATCACTTTGTCGCTGGGCTC |
| | Seq_SIC1_F | CATTGGGTCGTGTAAATAGG |
| | Seq_SIC1_R | CTGAGTGACCAGTTCATCTG |

#### Microfluidics and live cell imaging

Yeast cells were cultivated and imaged in custom-made microfluidic devices for all time-lapse microscopy experiments described in this study. One day before each experiment a new microfluidic chip was made by casting a mixture of 10 g of polydimethylsiloxane (PDMS; Sylgard 184 kit, Neyco) and 1 g of curing agent on a master wafer made by soft lithography. The chip was then degased, cured at 65°C for 4 hours, peeled off, punched with a 1.2 mm-diameter needle at all positions of inlets and outlets and then bonded onto a 24 x 60 mm coverslip after plasma activation of the surfaces. The chip pattern is shown on **Figure 1 - figure supplement 1a**. In brief, it consists of five independent pairs of flow channels (800 μm wide x 50 μm high) connected each to five growth chambers (400 x 400 x 3.8 μm; L x W x H) where yeast cells are constrained to proliferate in monolayer.

Yeast strains were thawed from glycerol stocks kept at -80°C onto YPG agar plates (10 g/L Yeast extract, 20g/L peptone, 5% (v/v) glycerol, 20 g/L bacto agar) at least three days and no more than two weeks before each microscopy experiment. After two days of incubation at 30°C, YPG plates were kept at room temperature. The day before an experiment, a small amount of cells (~$10^5$ cells) was inoculated in 4 ml of CSM medium (6.7 g/L Yeast Nitrogen Base without amino acids (BD Difco™), 2% glucose (Euromedex), 0,8 g/L Complete Supplement Mixture (CSM) of amino acids (MP Biomedicals™)) and incubated overnight at 30°C with 250 rpm orbital shaking. 100 μl of cell culture was then transferred to 5 ml of CSM medium and incubated for another 4 hours at 30°C to reach the exponential phase of growth. Cells were loaded in the microfluidic chip by injecting 50 μl of cell culture through each inlet of the chip using a small syringe.

Next, each pair of inlets was connected to a three-way solenoid valve (the Lee Company, ref. LFAA1201418H) that was itself connected to two bottles of medium via tubings with 0.5 mm inner diameter and 1.5 mm outer diameter (Cole-Parmer Microbore Tubing). A custom-made valve controller piloted by a Node-RED application was used to dynamically control which medium was dispensed to the cells based on a predetermined schedule of valve state switching. Media were sterilized by filtration (0.22 μm) to remove large particles and their composition varied depending on the experiment. All media were based on 6.7 g/L Yeast Nitrogen Base without amino acids (BD Difco™) and 0,8 g/L Complete Supplement Mixture (CSM) of amino acids (MP Biomedicals™) in $ddH_20$. This base was complemented either with 2% glucose, 0.1% glucose or no glucose and with 1 M sorbitol or no sorbitol. The outlets of the microfluidic chip were connected via tubings to a peristaltic pump (Ismatec IPC 12) and to an empty beaker to collect the flow through.

The chip was then mounted on a motorized inverted microscope (Olympus IX83) equipped with LEDs for fluorescence excitation (CoolLED pE-300ultra), a Zyla 4.2 sCMOS camera (Andor - Oxford Instruments) and an autofocus module (IX3-ZDC2, Olympus). The microscope, the microfluidic system and culture media were all placed inside an incubation chamber maintained at 30°C. Immediately after mounting the chip, the pump was turned on with a flow rate set at 120 μl/min for each outlet tube, resulting in a flow rate of 240 μl/min in the growth chambers since they were each connected to two outlets. The medium dispensed to cells by default was CSM with either 2% glucose or 0.1% glucose. The microscope was controlled using iQ v3.6.3 software

(Andor Technology) and all images were obtained using a 60x oil immersion objective (Olympus PlanApo N 60X/1.42) and a 1.6x magnification changer. Using live bright field imaging, we selected 25 positions (25 fields of view) of the motorized stage (Prior Scientific ProScan III) that captured 10 to 50 cells in each of the 25 growth chambers of the chip and were focused slightly below the median cell plane based on cell wall contrast. We used an iQ program that automatically scanned each position every 6 minutes for 12 or 24 hours and acquired bright field and fluorescence images after autofocus adjustment. In parallel, we executed the program controlling the timing of valve state switches. This program always started with one hour of the default state corresponding to CSM + 2% glucose or CSM + 0.1% glucose so that cells could acclimate to the experimental settings and images recorded in the first hour were excluded from analyses. To detect YFP fluorescence, samples were exposed to blue LED illumination at an intensity of 10% using a 514/10 nm excitation filter and fluorescence was acquired with an exposure time of 250 ms using a 545/40 nm emission filter. To detect mCherry fluorescence, samples were exposed to green LED illumination at an intensity of 14% using a 560/40 nm excitation filter and fluorescence was acquired with an exposure time of 150 ms using a 630/75 nm emission filter. Microscopy images were saved in TIFF format and are available upon request.

Microscopy images were acquired following the same procedure for experiments aiming at quantifying division and death rates, for experiments aiming at quantifying the fluorescence of cells expressing *P$_{STL1}$-mCitrine* and for experiments aiming at quantifying the nuclear enrichment of Hog1-GFP fusion protein. However, image analysis was performed differently for each type of experiment as described below.

#### Quantification of cell division and death rates

We used *ilastik* v1.3 for the segmentation and tracking of cell nuclei expressing HTB2-mCherry on fluorescence images. Three time-lapse movies of 120 fluorescence images of yPH_132 cells grown in CSM + 2% glucose medium were used to train the machine learning algorithms to recognize the nuclei of single cells and the nuclei of dividing cells at various numbers of cells per image. The segmentation and tracking procedure was then applied to frames from all experiments after manually removing out-of-focus images that occasionally occurred due to autofocus failure. The output for each experiment was a CSV file for each of the 25 field of views where rows corresponded to all objects (i.e. cell nuclei) detected on all images and columns corresponded to parameters such as image identity, cell nucleus identity, parental nucleus identity, size of the nucleus and XY coordinates of nucleus centroid. For all experiments involving combined stresses and alternating stresses (**Figures 3-6**), we manually added to CSV files a parameter indicating when cell death was observed based on visual inspection of bright field images. This manual step was necessary because the nucleus of a dead cell could remain fluorescent and could be tracked for several hours after cell death. This time-consuming step was not performed in the analyses of other experiments where death events remained very rare (results shown in **Figure 2**). Next, we computed cell division rate and death rate using homemade R scripts available in **Supplementary File 1**. We designed and implemented an Eulerian measure of division rate that was robust to rare tracking errors and to cell saturation in the field of view. In this approach, we first define a tracking window of 1928 x 1928 pixels centered on each image of 2048 x 2048 pixels. We can then establish the relation:

$$N_{window_{(t+1)}} - N_{window_t} = N_{new_{(t)}} + N_{in_{(t)}} - N_{out_{(t)}}$$

where $N_{window_{(t)}}$ and $N_{window_{(t+1)}}$ are the number of cell nuclei in the tracking window at frames $t$ and $t + 1$ , $N_{new_{(t)}}$ is the number of divisions that occurred in the window between $t$ and $t + 1$ , $N_{in_{(t)}}$ is the number of cell nuclei that entered the window and $N_{out_{(t)}}$ is the number of cell nuclei that exited the window between $t$ and $t + 1$ . From this relation, we computed $\frac{N_{new_{(t)}}}{N_{w_{(t)}}}$ which is the number of division events relative to the total number of cell nuclei in the window between $t$ and $t + 1$ . The slope of the linear regression of the cumulative sum $\sum_{t=t_0}^{t_1} \frac{N_{new_{(t)}}}{N_{window_{(t)}}}$ $t_0$ and $t_1$ . When death events were added to the input file, we only counted the nuclei of living cells when calculating the division rate. We excluded frames at the end of experiments that corresponded to an incomplete period of environmental fluctuation. The average death rate was calculated using a similar approach: it corresponded to the slope of the linear regression of the cumulative sum $\sum_{t=t_0}^{t_1} \frac{N_{death_{(t)}}}{N_{window_{(t)}}}$ time where $N_{death_{(t)}}$ was the number of death events observed in the tracking window between $t$ and $t + 1$ .

#### Quantification of *P<sub>STL1</sub>-mCitrine* expression

We used a custom image analysis approach based on the segmentation and tracking of cells in bright field images to quantify *P*$_{STL1}$*-mCitrine* expression in single cells (yPH_091 strain) over time. First, a preprocessing step was necessary to eliminate out-of-focus images that occasionally occurred. This was done using a U-NET model trained to estimate the area of cells on each frame and to detect sudden changes of cell area between consecutive frames indicative of autofocus failure. Next, a proper cell segmentation was achieved via a machine-learning algorithm based on the StarDist method (https://github.com/stardist/stardist) that uses star-convex shape prior. This method was well suited to the round shape of yeast cells and performed slightly better than the U-NET network allowing more reliable tracking of cells. Training of StarDist and U-NET algorithms was performed using a GPU NVIDIA RTX Quadro. For each cell in a frame $F_t$ , the tracking was performed by detecting which cell in the previous frame $F_{t-1}$ was closest to the cell on frame $F_t$ based on euclidean distances between cell centroids. This simple method worked best for cells that did not move too much between consecutive frames: it could fail when the distance between two different cells in two consecutive frames was smaller than the distance between the same cell in the two frames. For this reason, we manually selected by visual inspection cells with correct tracking over at least 6 hours and excluded cells with wrong tracking in further analyses. We computed the mean fluorescence of each cell as the total fluorescence of the cell divided by the area of the cell. The area corresponded to the number of pixels classified as belonging to the cell in the segmented bright field image. The total fluorescence of a cell was the sum of intensities of all pixels classified as belonging to the cell in the fluorescence image. Pixel classification of fluorescence images was performed using a *numpy.array* function in Python that applied the segmentation masks obtained from the analysis of bright field images to the corresponding fluorescence images. R scripts were used to plot the mean fold change of fluorescence over time for all cells analyzed in images that were taken at the same time point in different growth chambers sharing the same regime of environmental fluctuations. Fold change was calculated for each cell as the difference of mean fluorescence observed for that cell in a given frame and the mean fluorescence observed among all cells in the first frame divided by the mean fluorescence among all cells in the first frame.

#### Hog1-GFP nuclear enrichment

The nuclear enrichment of Hog1-GFP was quantified in cells expressing both the Hog1-GFP reporter and the nuclear marker Htb2-mCherry. Cell segmentation and tracking was performed on bright field images following the same procedure as described above for the quantification of *P<sub>STL1</sub>-mCitrine* expression. In addition, another segmentation was done for cell nuclei that were detected with the red fluorescence channel using a simple thresholding step. Each nucleus contour was then associated by contour overlapping comparison to its corresponding cell contour obtained by segmentation of bright field images. We then computed the mean fluorescence in the green channel for each cell and for each nucleus. The mean fluorescence of a cell was calculated as the total intensity of all pixels classified as belonging to the cell (including the nucleus) divided by the number of these pixels. The mean fluorescence of a nucleus was calculated as the total intensity of all pixels classified as belonging to the nucleus divided by the number of these pixels. Finally, the nuclear enrichment of fluorescence was calculated for each cell as the mean fluorescence of the nucleus divided by the mean fluorescence of the cell. R scripts were used to plot the nuclear enrichment of Hog1-GFP fluorescence over time for all cells analyzed in images that were taken at the same time point in different growth chambers sharing the same regime of environmental fluctuations.

#### Fluorescein assay

We used a fluorescein assay to characterize the temporal dynamics of medium fluctuations inside microfluidic chips. In this assay, we connected a microfluidic chip to CSM medium complemented with 50 nM fluorescein and to CSM medium without fluorescein. We programmed the valve to dispense CSM with fluorescein to the chip for 20 minutes followed by CSM without fluorescein for another 20 minutes and repeated this treatment twice. We tried different flow rates on the peristaltic pump but only showed results for the optimal flow rate of 120 µl/min. We used the same microscopy setup as described above to image the growth chamber at the center of the chip, except that a 20x objective (Olympus Plan Achromat) was used to be able to visualize both the flow channel and the growth chamber in the field of view. One bright field image and one fluorescence image were taken every 12 seconds. The fluorescence channel consisted of blue LED illumination at an intensity of 10% with a 514/10 nm excitation filter and acquisition with an exposure time of 250 ms using a 545/40 nm emission filter. We used Image J to quantify the mean fluorescence in two circular regions with a diameter of 280 pixels. One region was in the center of the growth chamber and the other region in the flow channel. A script in R was used to plot the relative level of fluorescence over time in each region. The relative fluorescence at time $t$ ($RF_t$) was calculated as $RF_t = \frac{F_t - F_{min}}{F_{max} - F_{min}}$ where $F_t$ is the mean fluorescence at time $t$, $F_{min}$ is the mean fluorescence observed between 30 and 40 minutes and between 70 and 80 minutes when fluorescein was at its minimal concentration in the chip and $F_{max}$ is the mean fluorescence observed between 10 and 20 minutes and between 50 and 60 minutes when fluorescein was at its maximal concentration in the chip.

### Figure 1 – figure supplement 1

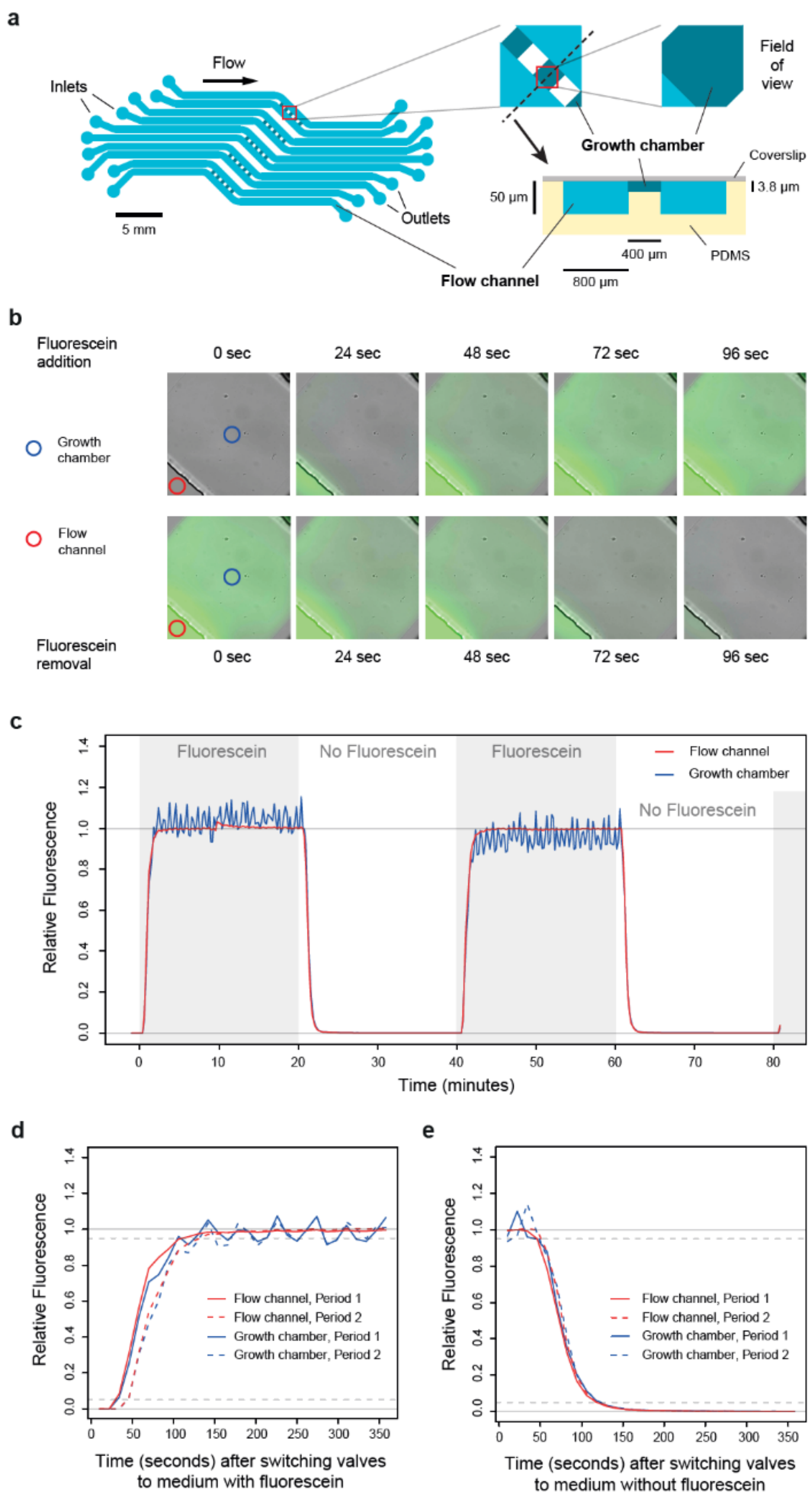


**Caption Figure 1 – figure supplement 1. Dynamics of media switching inside the microfluidic chip. (a)** Design of the microfluidic device used in this study. Cells are trapped in five sets of five growth chambers (400 x 400 x 3.8 μm each) located in the center of the PDMS chip. Each set of five growth chambers is connected to two flow channels to allow rapid diffusion of the medium inside the chambers. The dynamics and composition of media can be independently controlled for each set. The pattern of the microfluidic device (at scale) is shown on the left, with a close up caption of a growth chamber shown on the top right. A transverse view (out of scale) along the black dotted line is shown on the bottom right. **(b)** Images showing fluorescence dynamics in a growth chamber after fluorescein (50 nM) is added (top row) or removed (bottom row) from the medium. Blue and red circles show the two positions where fluorescence was quantified in **c-e.** **(c)** Quantification of fluorescence dynamics during periodic switches between SC medium with and without fluorescein. **(d)** Relative fluorescence measured during 350 seconds after switching the valve from medium without fluorescein to medium with fluorescein. **(e)** Relative fluorescence measured during 350 seconds after switching the valve from medium with fluorescein to medium without fluorescein. Data shown in **(d)** and **(e)** are also shown in **(c)** (same experiment). **(c-e)** Fluorescence was quantified both in the flow channel (red line) and in the middle of the growth chamber (blue line). Fluorescence is expressed on a relative scale (see Methods) to focus the comparison on the temporal dynamics instead of the absolute fluorescence level (the fluorescence intensity is much lower in the growth chamber because it is thinner than the flow channel). Acquisition of bright field and fluorescence images was performed once every 12 seconds.

### Figure 2 – figure supplement 1

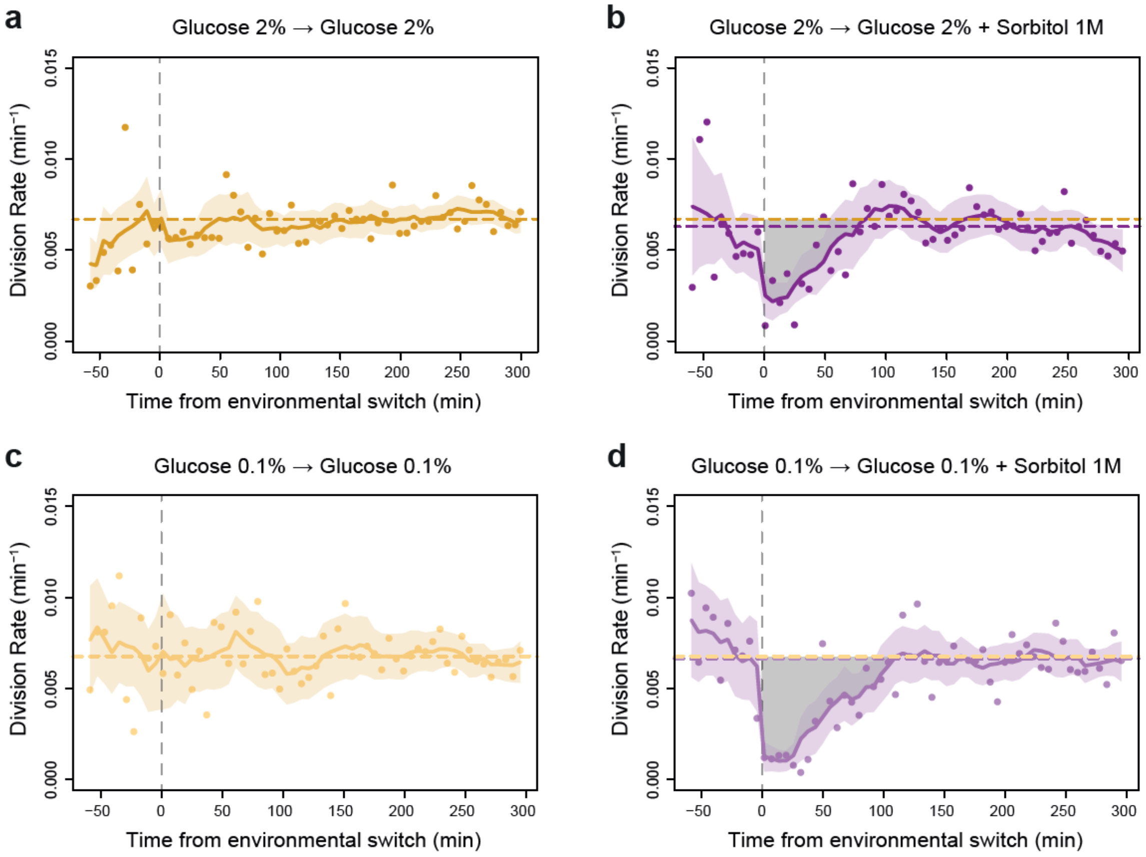


**Caption Figure 2 – figure supplement 1. Impact of a single hyperosmotic shock on cell division rate. (a-d)** Temporal dynamics of division rate as shown in **Figure 2d** but with confidence intervals and individual data points. Colored lines are the same as in **Figure 2d**. Colored areas represent 95% confidence intervals of the average division rate. Colored dots show the average division rate measured every 6 minutes (not averaged across sliding windows) among all growth chambers exposed to the same medium. Horizontal dotted lines show the average division rate measured between 100 and 300 minutes after medium switching (i.e., after adaptation to the new medium) in the absence (orange) or presence (purple) of osmotic stress. Vertical dotted lines show time $t = 0$ when the medium was changed in the microfluidic chip. **(b,d)** Gray areas represent the cost in cell divisions of one sustained hyperosmotic shock (1 M sorbitol) in **(b)** 2% glucose or **(d)** 0.1% glucose. The initial number of cells analyzed among replicates ranged from 220 to 776.

### Figure 2 – figure supplement 2

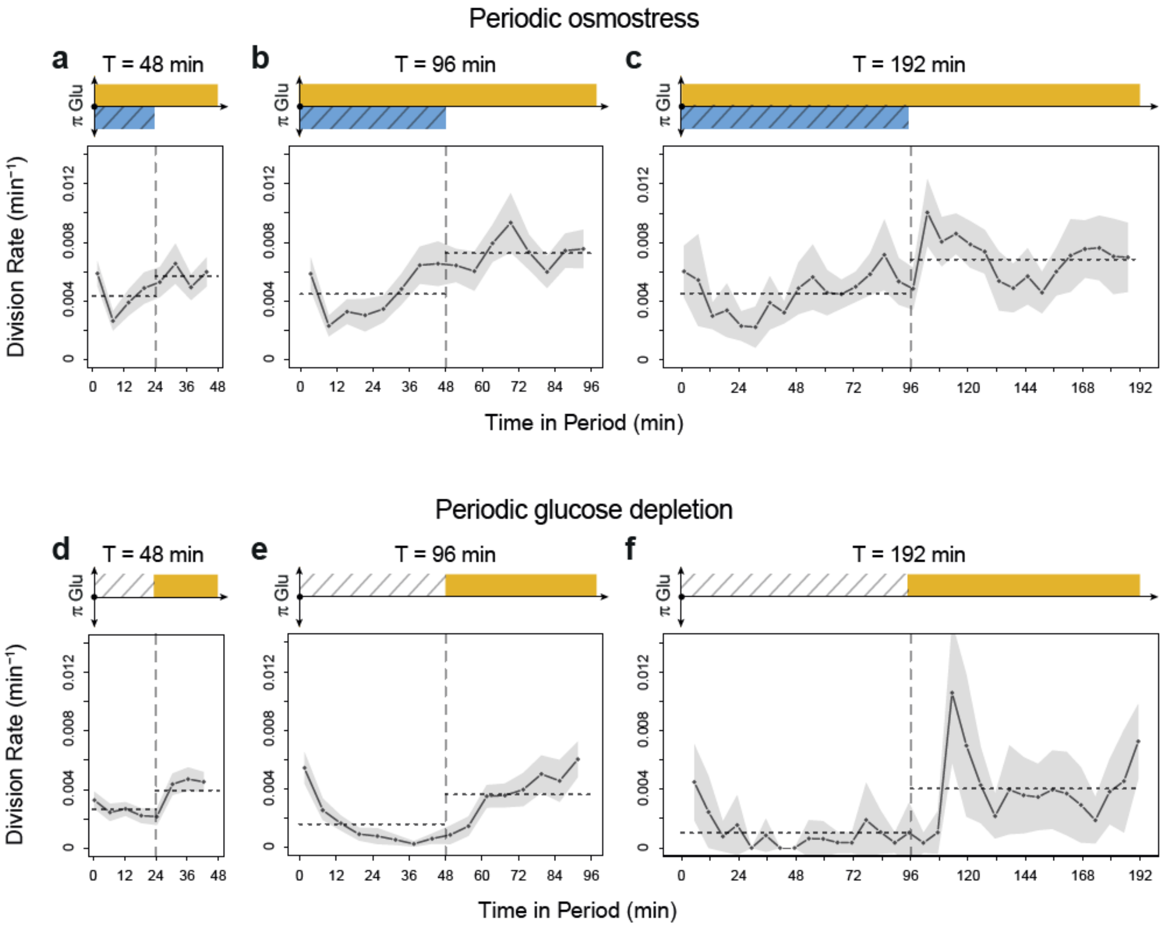


**Caption Figure 2 – figure supplement 3. Temporal dynamics of division rate under periodic osmotic stress and under periodic glucose depletion.** In each plot, division rates measured over several consecutive periods of environmental fluctuations are averaged in a single period: each dot shows the average division rate during a 6-minute window centered on that dot for all fields of view sharing the same condition and for all periods in the experiment. **(a-c)** Temporal dynamics of division rate during fluctuations of osmotic stress (1M Sorbitol) with periods of 48 minutes **(a)**, 96 minutes **(b)** and 192 minutes **(c)**. **(d-f)** Temporal dynamics of division rate during glucose fluctuations (from 2% glucose to 0% glucose) with periods of 48 minutes **(d)**, 96 minutes **(e)** and 192 minutes **(f)**. Gray areas are 95% confidence intervals of the mean division rate. Horizontal dotted lines show the mean division rate for all data collected in each half period. The colored bars represent the periodic fluctuations of glucose (orange) and/or sorbitol (blue); hatching represents stress.

### Figure 2 – figure supplement 3

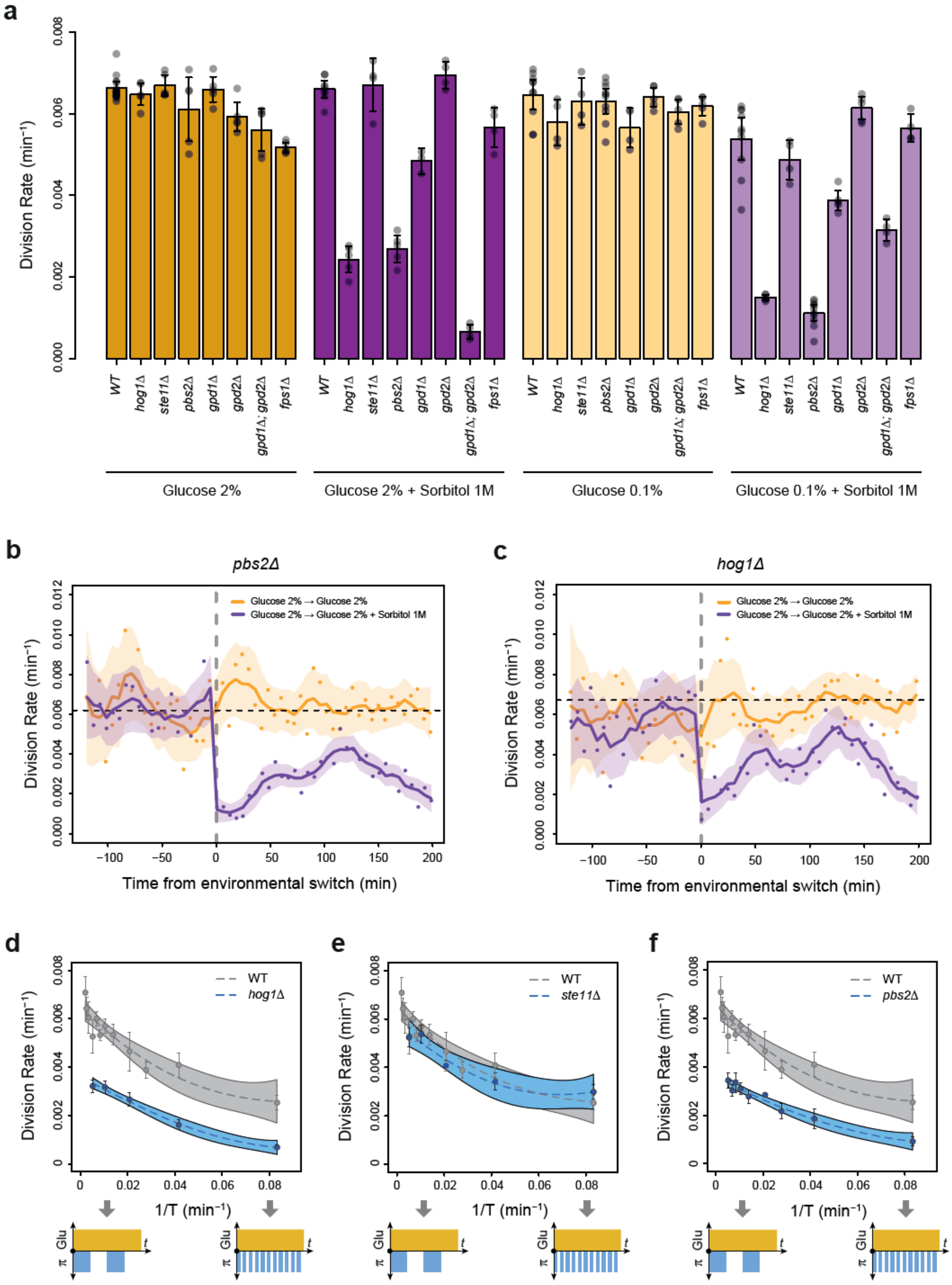

**Caption Figure 2 – figure supplement 3. Division rates of HOG pathway mutants under constant and periodic osmotic stress. (a)** Division rates of the wild type strain and of seven deletion mutants in four steady conditions. Bars show the mean division rate measured among growth chambers sharing the same environmental condition. Error bars are 95% confidence intervals of the mean. Dark dots show the average division rate for each field of view. Colors represent the different environmental conditions indicated at the bottom. **(b,c)** Temporal dynamics of division rate of **(b)** *pbs2Δ* mutant cells and **(c)** *hog1Δ* mutant cells under sustained hyperosmotic stress (purple line, 1 M sorbitol added at $t = 0$ min) or in standard condition with 2% glucose (orange line). Each curve represents the "instantaneous" division rate calculated every 6 minutes across sliding windows of 36 minutes (see Methods) and averaged for cells imaged at several positions in the microfluidic chip. Colored areas represent 95% confidence intervals of the average division rate. Colored dots show the average division rate measured every 6 minutes (not averaged across sliding windows) among all growth chambers exposed to the same medium. Horizontal dotted lines show the average division rate measured between 100 and 300 minutes after medium switching (i.e., after adaptation to the new medium) in the absence of osmotic stress. Vertical dotted lines show the time $t = 0$ when the medium was changed in the microfluidic chip. **(d-f)** Relationship between the frequency of osmotic stress and division rate for **(d)** *hog1Δ*, **(e)** *ste11Δ* and **(f)** *pbs2Δ* mutant cells. Data collected for each mutant strain are shown in blue, while data collected for the wild type strain (yPH_132) are shown in gray as reference. Each dot shows the mean division rate measured among different growth chambers exposed to the same condition. Error bars are 95% confidence intervals of the mean. Dotted lines are Loess regressions obtained with a smoothing parameter of 0.66. Filled areas represent 95% confidence intervals of the regression estimates. The initial number of cells analyzed among replicates ranged from 77 to 1753 in **(a)**, from 75 to 386 in **(d)**, from 75 to 539 in **(e)** and from 53 to 386 in **(f)**.

### Figure 3 – figure supplement 1

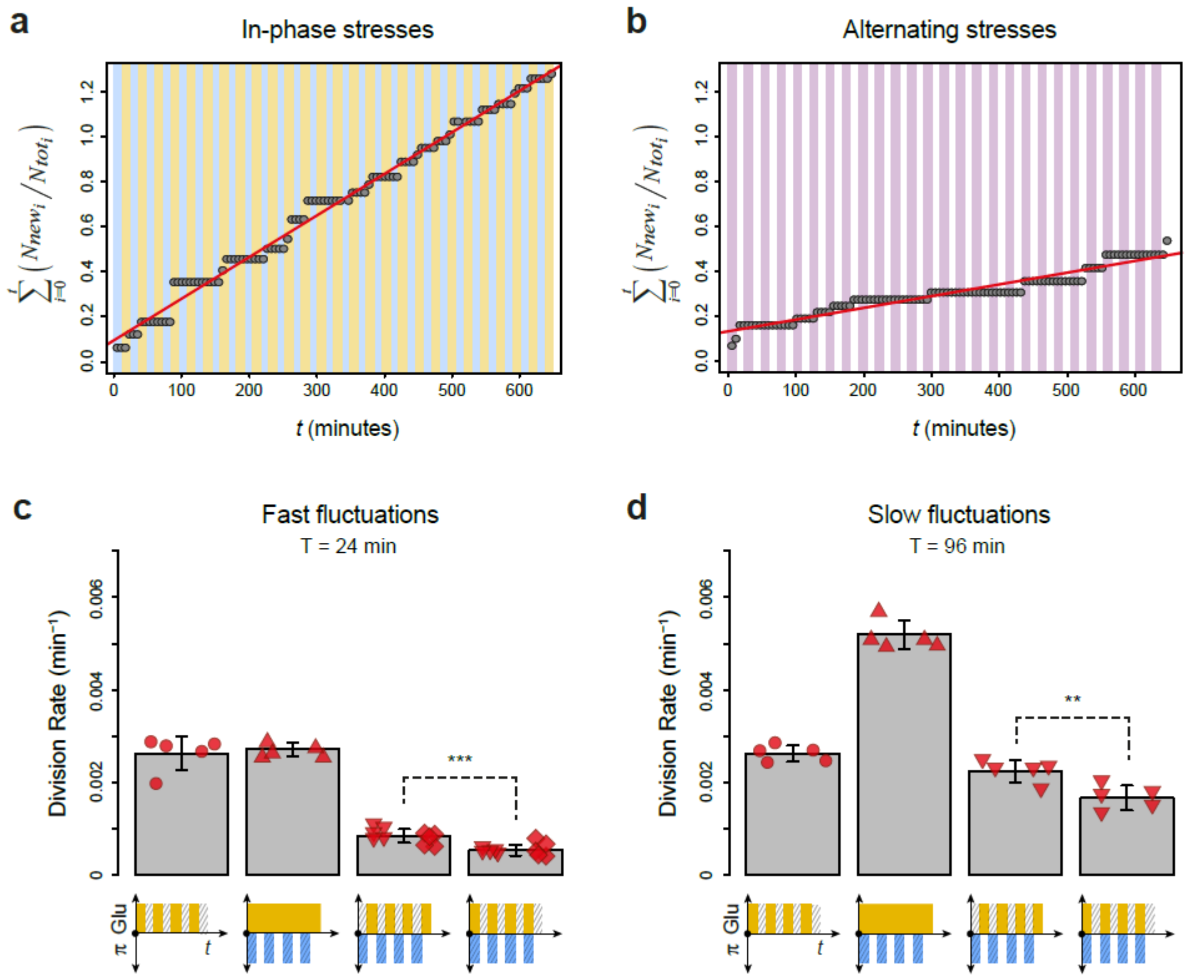


**Caption Figure 3 – figure supplement 1**. **Division rates under regimes of in-phase stresses and alternating stresses with different fluctuation periods and glucose concentrations. (a)** Accumulation of new cells as a function of time under IPS condition (periodic growth in medium with 2% glucose for 12 minutes, followed by medium without glucose and with 1 M sorbitol for 12 minutes). **(b)** Accumulation of new cells as a function of time under AS condition (periodic growth in medium with 2% glucose and 1 M sorbitol for 12 minutes, followed medium without glucose or sorbitol for 12 minutes). **(a,b)** Dots show the cumulative number of new cells over the total number of cells in one field of view as a function of time. The average cell division rate was calculated at the slope of the regression line (red line). Vertical colored bars show the timing of media switches (orange: 2% glucose and no sorbitol; blue: no glucose + 1 M sorbitol; purple: 2% glucose + 1 M sorbitol; white: no glucose and no sorbitol). **(c,d)** Mean cell division rate under fluctuations of 1M sorbitol and/or 0.1% glucose with a period of **(c)** 24 minutes or **(d)** 96 minutes. The four conditions are periodic glucose deprivation, periodic osmostress, in-phase stresses and alternating stresses. Bars represent the mean division rate among different growth chambers. Error bars are 95% confidence intervals of the mean. Red symbols show the mean division rate for each growth chamber (field of view), with different symbols representing data from experiments performed on different days with different microfluidic chips. *t*-tests were performed to compare the mean division rate between in-phase stresses and alternating stresses conditions (*** : $P < 0.001$; ** : $0.001 \leq P < 0.01$). Colored graphs are used to represent the periodic fluctuations of glucose (orange) and/or sorbitol (blue) in the medium; hatching represents stress. The initial number of cells analyzed among replicates ranged from 149 to 306 in **(c)** and from 83 to 136 in **(d)**.

### Figure 4 – figure supplement 1

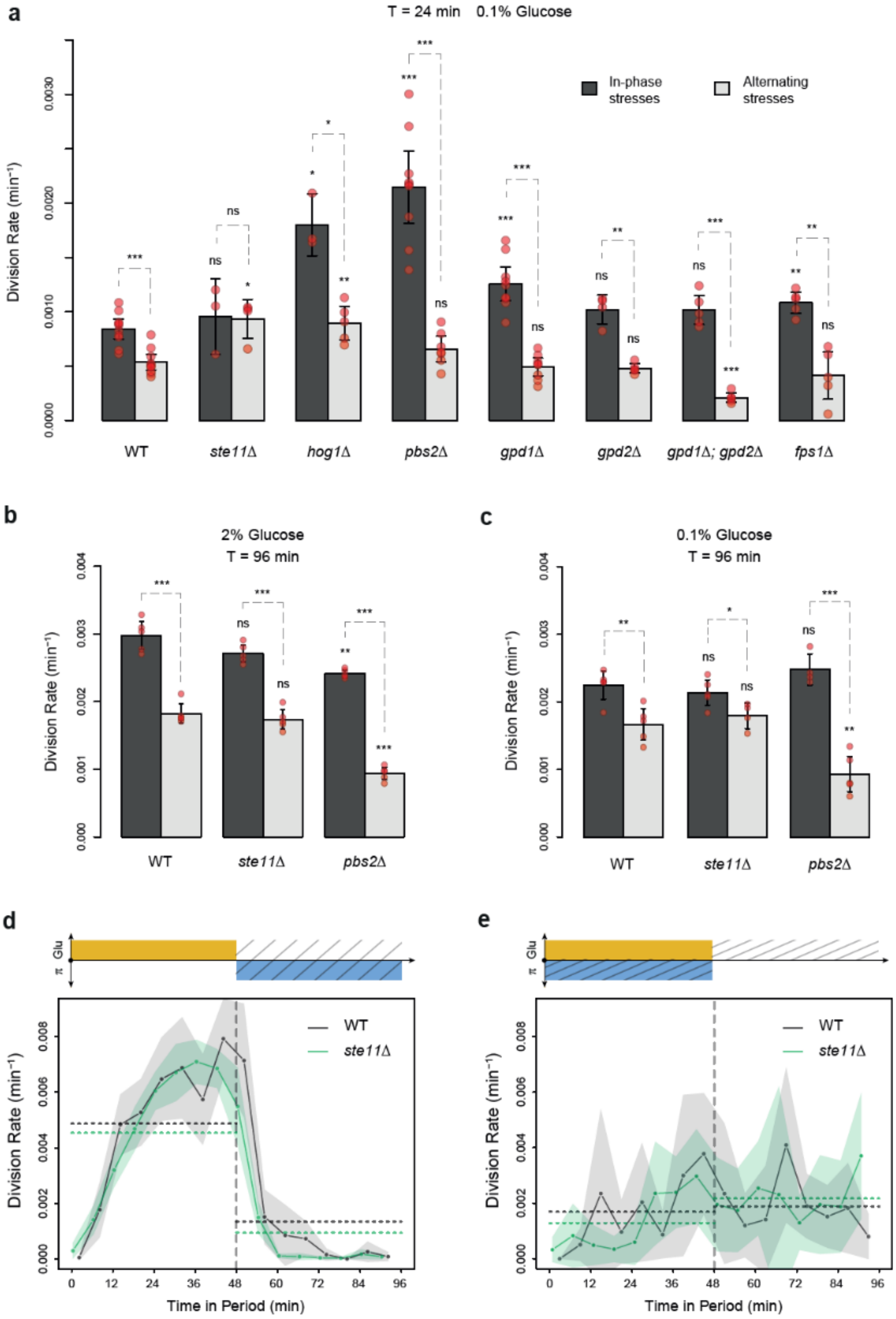


**Caption Figure 4 – figure supplement 1. Division rates of HOG pathway mutants during in-phase stresses and alternating stresses with different fluctuation periods and glucose**

**concentrations. (a-c)** Division rate of deletion mutants during growth in IPS and AS conditions with **(a)** a period of 24 minutes and 0.1% glucose, **(b)** a period of 96 minutes and 2% glucose and **(c)** a period of 96 minutes and 0.1% glucose. Bars show the mean division rate among growth chambers sharing the same environmental conditions. Error bars are 95% confidence intervals of the mean. Red symbols show the average division rate for each growth chamber. Results of *t*-tests comparing the wild-type and mutant strains under the same conditions are indicated above each bar; results comparing the same strain under different conditions are shown above each pair of bars (ns: $P > 0.05$; * $0.01 < P < 0.05$; ** $0.001 < P < 0.01$; *** $P < 0.001$). **(d-e)** Temporal dynamics of division rate during a period of 96 minutes in **(b)** IPS and **(c)** AS conditions for wild-type (black) and *ste11Δ* mutant (green) cells. Each dot shows the division rate during a 6-minute window centered on that dot and averaged for all fields of view sharing the same condition and all periods in the experiment. Gray and green areas are 95% confidence intervals of the mean division rate. Horizontal dotted lines show the mean division rate for all data collected in each half period. The colored bars represent the periodic fluctuations in glucose (orange) and/or sorbitol (blue); hatching represents stress. The initial number of cells analyzed among replicates ranged from 22 to 411 in **(a),** from 94 to 626 in **(b)**, and from 67 to 206 in **(c)**.

### Figure 4 – figure supplement 2

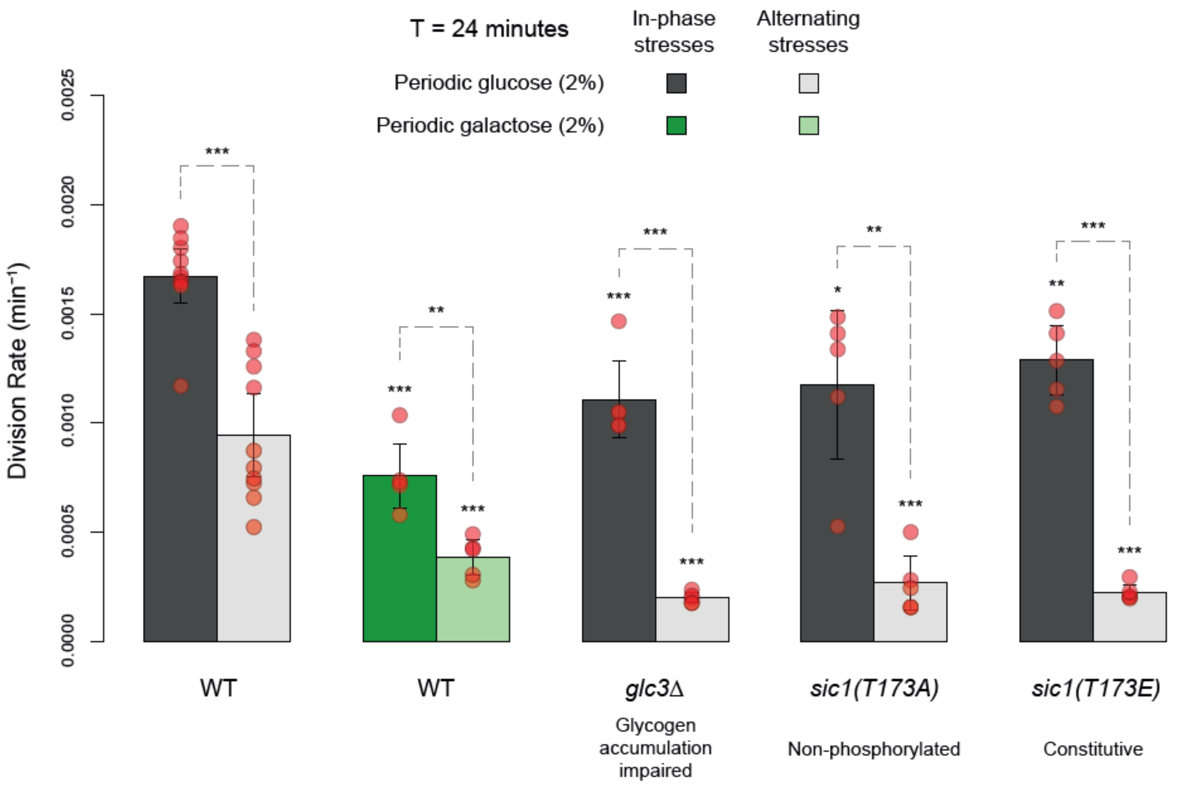


**Caption Figure 4 – figure supplement 2. Comparing division rates during in-phase stresses and alternating stresses of wild-type cells exposed to periodic glucose (gray) or periodic galactose (green) and of mutant cells with impaired glycogen accumulation (*glc3* deletion) or impaired cell cycle arrest in response to hyperosmotic stress (*sic1* alleles).** The period of environmental fluctuations is 24 minutes in all conditions. Bars show the mean division rate among growth chambers sharing the same environmental conditions. Error bars are 95% confidence intervals of the mean. Red symbols show the average division rate for each growth chamber. Results of *t*-tests comparing the wild-type and mutant strains under the same conditions are indicated above each bar; results comparing the same strain under different conditions are shown above each pair of bars (ns: $P > 0.05$; * $0.01 < P < 0.05$; ** $0.001 < P < 0.01$; *** $P < 0.001$). The initial number of cells analyzed among replicates ranged from 14 to 120.

### Figure 5 – figure supplement 1

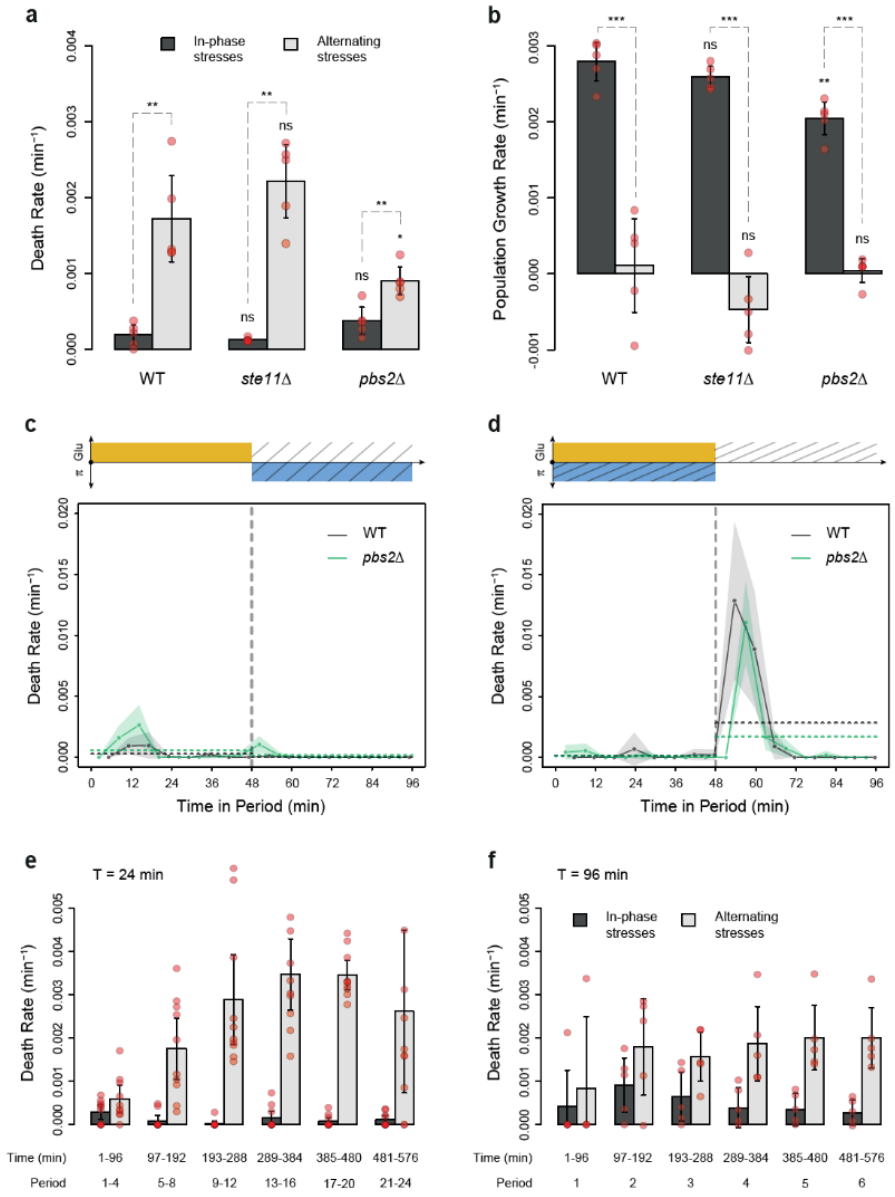


**Figure 5 – figure supplement 1**. **Dynamics of cell death and population growth rates under combined and alternating stresses. (a)** Death rates and **(b)** population growth rates (calculated as the difference between the division rate and death rate) in IPS (dark gray) and AS (light gray) conditions with a period of 96 minutes and 2% glucose. **(a,b)** Bars show the mean values measured among different growth chambers sharing the same environmental condition. Error bars are 95% confidence intervals of the mean. Red symbols show the average death rate or population growth rate for each growth chamber. Results of *t*-tests comparing the wild-type and mutant strains under the same conditions are indicated above each bar; results comparing the same strain under different conditions are shown above each pair of bars (ns, $P > 0.05$; * $0.01 < P < 0.05$; ** $0.001 < P < 0.01$; *** $P < 0.001$). **(c,d)** Temporal dynamics of death rate averaged across all

96-minute periods of growth in **(c)** IPS and **(d)** AS conditions for wild type (black) and *pbs2Δ* mutant (green) cells. Each dot shows the death rate during a window of 6 minutes centered on that dot and averaged across all fields of view sharing the same conditions and among all periods in the experiment. Gray and green areas are 95% confidence intervals of the mean division rate. Horizontal dotted lines show the mean death rate among all data collected in each half period (*i.e.,* in each medium since the medium is changed at the start and the middle of each period (vertical dotted line)). Colored graphs are used to represent the periodic fluctuations of glucose (orange) and/or sorbitol (blue) in the medium; hatching represents stress. **(e,f)** Average death rates measured at six intervals of time in IPS (dark gray) and AS (light gray) conditions at a period of **(e)** 24 minutes or **(f)** 96 minutes. Bars show the mean death rate measured in different growth chambers sharing the same conditions. Error bars are 95% confidence intervals of the mean. Red symbols show the average death rate for each field of view and time interval.

### Figure 5 – figure supplement 2

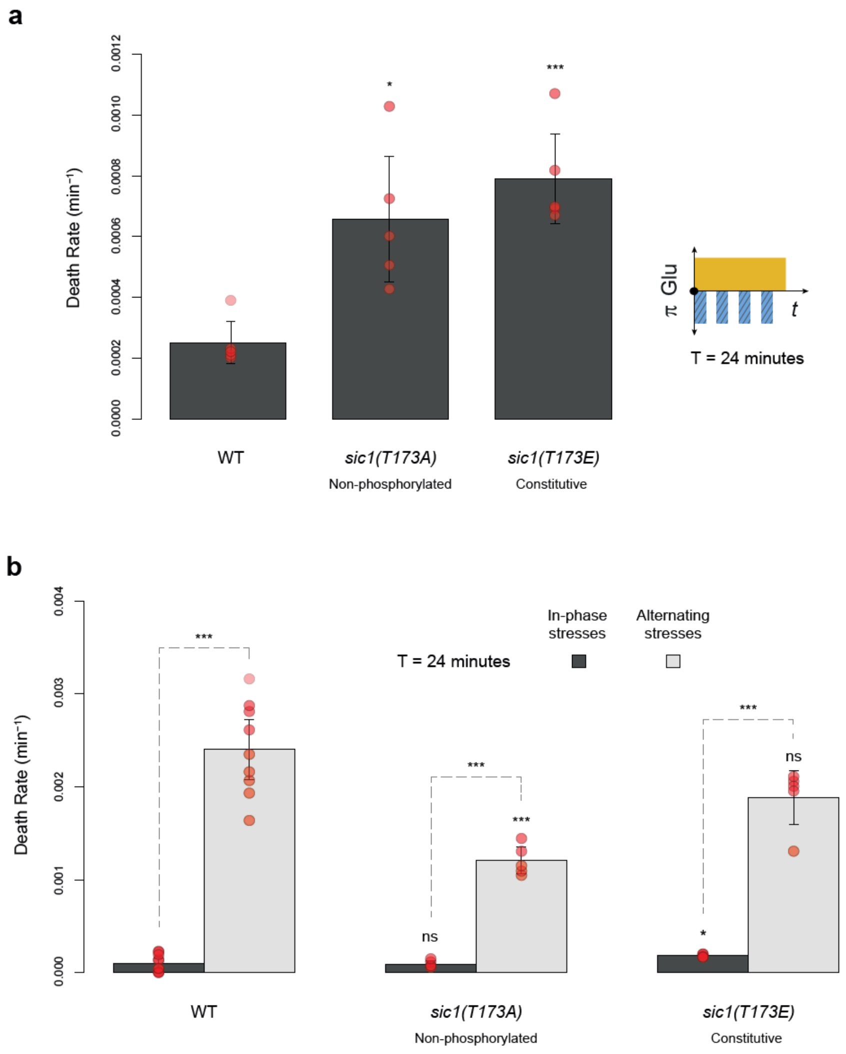


**Figure 5 – figure supplement 2. Death rates of wild-type and *sic1* mutant cells under periodic fluctuations of hyperosmotic stress with or without fluctuations of glucose availability. (a)** Death rates in constant 2% glucose with repeated addition and removal of 1M sorbitol at a period of 24 minutes. **(b)** Death rates in IPS (dark gray) and AS (light gray) conditions with a period of 24 minutes and 2% glucose. **(a,b)** Bars show the mean death rates measured among different growth chambers sharing the same environmental condition. Error bars are 95% confidence intervals of the mean. Red symbols show the average death rate for each growth chamber. Results of *t*-tests comparing the wild-type and mutant strains under the same conditions are indicated above bars (ns, $P > 0.05$; * $0.01 < P < 0.05$; ** $0.001 < P < 0.01$; *** $P < 0.001$). The initial number of cells analyzed among replicates ranged from 14 to 120.

### Figure 6 – figure supplement 1

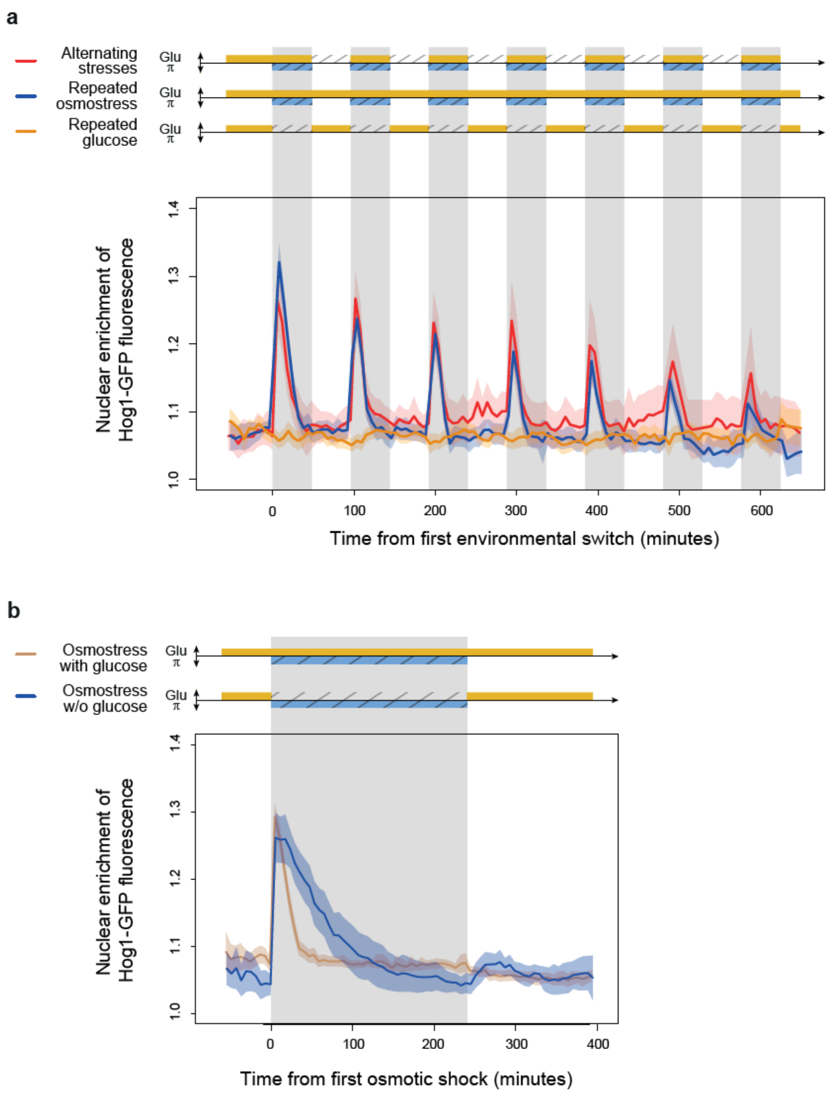


**Caption Figure 6 – figure supplement 1**. **Temporal dynamics of Hog1-GFP location under different regimes of environmental fluctuations. (a,b)** Nuclear enrichment of Hog1-GFP fluorescence over time under **(b)** alternating stresses (red), periodic osmotic stress in constant glucose (blue) and periodic glucose fluctuations in absence of osmotic stress or **(c)** a single osmotic shock in presence (brown) or absence (blue) of 2% glucose. Each curve shows the mean nuclear enrichment of GFP fluorescence measured for 11–97 cells from one to seven fields of view. Colored areas indicate 95% confidence intervals of the mean. The colored graphs represent the periodic fluctuations of 2% glucose (orange) and/or 1 M sorbitol (blue) in the medium; hatching represents stress.
